# Local 5-HT signal bi-directionally regulates the coincidence time window of associative learning

**DOI:** 10.1101/2022.03.27.485970

**Authors:** Jianzhi Zeng, Xuelin Li, Zimo Zhangren, Mingyue Lv, Yipan Wang, Ke Tan, Xiju Xia, Jinxia Wan, Miao Jing, Yang Yang, Yan Li, Yulong Li

**Author notes:** Manuscript correspondence: Yulong Li. These authors contributed equally: Jianzhi Zeng, Xuelin Li.

## Abstract

Temporal coincidence between the conditioned stimulus (CS) and unconditioned stimulus (US) is essential for associative learning across species. Despite its ubiquitous presence, the mechanism that may regulate this time window duration remains unclear yet. Using olfactory associative learning in *Drosophila* as a model, we find that suppressing or promoting serotonin (5-HT) signal could respectively shorten or prolong the coincidence time window of odor-shock associative learning and synaptic plasticity in mushroom body (MB) Kenyon cells (KCs). Capitalizing on <u>G</u>PC<u>R</u>-<u>a</u>ctivation <u>b</u>ased (GRAB) sensors for 5-HT and acetylcholine (ACh), we characterized the *in vivo* 5-HT dynamics in MB lobes during odor and shock stimulations and further dissected this microcircuit. Interestingly, local KC-released ACh activates nicotinic receptors on the dorsal paired medial (DPM) neuron, and in turn the DPM neuron releases 5-HT to inhibit the ACh signal via the 5-HT1a receptor. Finally, we demonstrated that the DPM-mediated serotonergic feedback circuit is sufficient and necessary to regulate the coincidence time window. This work provides a model for studying the temporal contingency of environmental events and their causal relationship.

## Main

To survive and proliferate in constantly changing environments, animals including humans have evolved associative learning to build a causal relationship between the neutral conditioned stimulus (CS) and the punitive or rewarding unconditioned stimulus (US). A prerequisite for successful associative learning is that the inter-stimulus interval (ISI) between two stimuli must fall within a relative short time window, also called the coincidence time window. The temporal contingency is critical for both Pavlovian conditioning (Pavlov and Anrep, 1927) and operant conditioning (Skinner, 1938) in a wide range of behaviors across species, including the siphon withdrawal reflex in *Aplysia* (Carew et al., 1981; Hawkins et al., 1986), olfactory associative learning in *Drosophila* (Tully and Quinn, 1985) and the eye-blinking task in humans (Bernstein, 1934; McAllister, 1953). Significantly, an altered coincidence time window has been associated with a variety of neurodevelopmental disorders, brain injuries, psychological diseases and psychedelic states (Bolbecker et al., 2011; Frings et al., 2010; Harvey, 2003; Harvey et al., 1988; McGlinchey-Berroth et al., 1999; Oristaglio et al., 2013; Perrett et al., 1993; Woodruff-Pak and Papka, 1996). Experimental evidences and computational theories have suggested that neuromodulatory signals play essential roles in the temporal discrimination of spike-timing-dependent plasticity (STDP), which is a cellular model for learning (Brzosko et al., 2019; Liu et al., 2020a; Pawlak et al., 2010). However, the underlying molecular or circuit basis for regulating the coincidence time window remains incompletely understood. Unraveling these mechanisms will provide valuable insights into how the brain determines the relationship between temporally discrete events and may shed new light on how brain disorders affect learning and memory.

Mushroom body (MB) is the major region involved in olfactory associative learning in *Drosophila*, which has highly ordered architecture and abundant genetic tools (Aso et al., 2014; Heisenberg, 2003; Mao and Davis, 2009; Tanaka et al., 2008), making it an ideal model for addressing fundamental questions regarding learning and memory. Recent progress in *Drosophila* brain electron microscopy (EM) connectomics (Eichler et al., 2017; Li et al., 2020; Scheffer et al., 2020; Takemura et al., 2017) and MB transcriptomics (Aso and Rubin, 2016; Croset et al., 2018) have provided additional evidences and will accelerate functional studies. The MB primarily consists of ∼2000 Kenyon cells (KCs) per hemisphere, with their dendrites forming the calyx and their axons bundled into three lobes, called the α/β lobe, α’/β’ lobe and γ lobe. These lobes are further segmented into 15 compartments, which are tiled by the axonal projections of dopaminergic neurons (DANs) and the corresponding dendrites arising from mushroom body output neurons (MBONs). During olfactory learning, KCs receive the CS signal from the olfactory circuit and punitive or rewarding US signal from DANs (Burke et al., 2012; Claridge-Chang et al., 2009; Kim et al., 2007; Liu et al., 2012; Qin et al., 2012; Schroll et al., 2006; Schwaerzel et al., 2003). Besides DA, other neuromodulators also converge on this MB microcircuit, including octopamine (OA), gamma-aminobutyric acid (GABA), 5-HT and glutamate.

The temporal relationship between the CS and US affects olfactory learning in *Drosophila* in two major aspects. First, the CS-US and US-CS pairing yield memories with opposite valence and this phenomenon is attributed to different dopamine receptors and intracellular cascades (Berry et al., 2012; Berry et al., 2018; Cohn et al., 2015; Handler et al., 2019; Hige et al., 2015; Himmelreich et al., 2017). Second, with a fixed temporal order such as CS-US pairing, the learning index declines as the interval between the CS and US increases, with a coincidence time window on the order of tens of seconds (Aso and Rubin, 2016; Gerber et al., 2019; Gerber et al., 2014; Tanimoto et al., 2004; Tomchik and Davis, 2009; Tully and Quinn, 1985). However, the specific neuromodulator and circuit-based mechanism that regulate the coincidence time window is currently unknown.

5-HT plays a critical role in learning and memory across species, including *Aplysia* (Kandel, 2001; Kandel and Schwartz, 1982), *C. elegans* (Zhang et al., 2005), mice (Fonseca et al., 2015; Li et al., 2016; Liu et al., 2014; Lottem et al., 2018; Miyazaki et al., 2018; Ren et al., 2018), humans (Buhot et al., 2000; Liu et al., 2020b) and *Drosophila*. The essential role of 5-HT in *Drosophila* learning and memory was firstly established in a place-learning paradigm (Sitaraman et al., 2008). In each hemisphere of the MB, the serotonergic DPM neuron innervates all three lobes, which has been reported to be involved in olfactory learning in both adults and larvae. (Ganguly et al., 2020; Johnson et al., 2011; Keene et al., 2006; Keene et al., 2004; Krashes et al., 2007; Lee et al., 2011; Waddell et al., 2000; Wu et al., 2011; Yu et al., 2005). However, the *in vivo* dynamics of 5-HT release from the DPM neuron, in responses to physiological stimuli and its regulation, are poorly understood. Moreover, little is known regarding how 5-HT affects the learning circuit in the MB.

In this work, we found that the coincidence time window for olfactory associative learning could be regulated by 5-HT in *Drosophila*. Taking advantage of the <u>G</u>PC<u>R a</u>ctivation‒<u>b</u>ased sensors for ACh (GRABACh3.0, ACh3.0) (Jing et al., 2020; Jing et al., 2018), we varied the CS-US the coincidence time window while monitoring KC-MBON synaptic plasticity, and found that it is regulated by 5-HT levels. Moreover, using GRAB5-HT1.0 (5-HT1.0) (Wan et al., 2021) we observed compartmental 5- HT signals in response to the odorant application and electric shock and identified the DPM neuron as the source of these 5-HT signals. Combining functional imaging with optogenetics and pharmacology, we found that the DPM neuron receives local excitation from KCs and then provides inhibitory serotonergic feedback to KCs. In addition, suppressing or promoting 5-HT release from DPM neurons respectively shortens or prolongs the coincidence time window of synaptic plasticity and learning behavior. These results suggest that the coincidence time window can be selectively regulated by local 5-HT release from DPM neurons in MB, which is critical for the organisms to efficiently form the correlation between environmental CS and US.

## Results

### 5-HT modulates the coincidence time window of one-trial olfactory learning behavior

To measure the coincidence time window of olfactory associative learning, we used the T-maze paradigm to train flies by pairing a 10-s odorant (CS+) and electric shocks (US) with varying inter-stimulus intervals (ISI), and presented another odorant (CS-) as an unpaired stimulus. After training, we tested flies’ performance index towards the CS+ and CS- (Figures 1A and 1B). We found that control flies (Canton-S) learned to avoid the CS+ when the ISI is ≤15 s, but had poor or no learning at longer ISI (Figure 1C). We used a sigmoid function to fit the relationship between the relative performance index against the ISI and the coincidence time window was indicated by the t50 of the fitted curve, which is 16.9 s for the control group. Next, we wanted to figure out whether the coincidence time window could be regulated by a specific neuromodulator, we focused on 5-HT due to its unclear function in short-term memory. By preventing 5-HT production through mutating the tryptophan hydroxylase (Trh) gene (Qian et al., 2017), which encodes the rate-limiting enzyme in 5-HT biosynthesis, we found that the coincidence time window was shortened to 10.8 s (Figure 1D). Given that the CS+ duration is 10 s, it means that Trh mutant flies cannot learn as soon as the CS and US cease to overlap. Conversely, when flies were pretreated with the selective serotonin reuptake inhibitor (SSRI) that is thought to elevate synaptic 5-HT levels (Ries et al., 2017; Yuan et al., 2005), the coincidence time window was extended to 25.2 s (Figure 1E). These results suggest that the coincidence time window in aversive associative learning can be bi-directionally regulated by the neuromodulator 5-HT.

**Figure 1.**
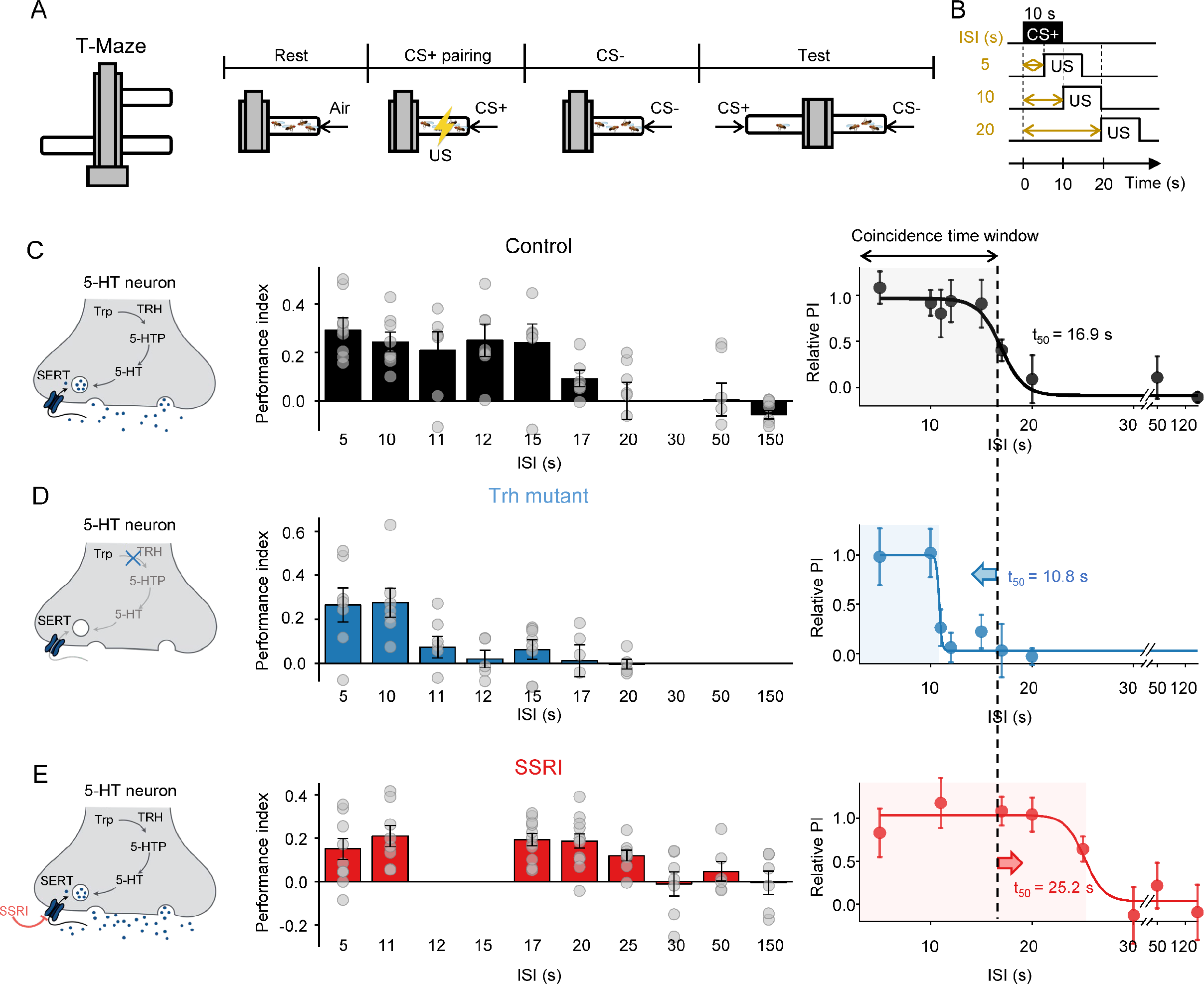
5-HT signaling can bi-directionally regulate the coincidence time window of olfactory learning. (**A**-**B**) Schematic diagram depicting the T-maze protocol (**A**) for measuring how the inter-stimulus interval (ISI) affects odorant-shock pairing-induced aversive memory (**B**). (**C-E**) Schematic diagram depicting the 5-HT synthesis process (left). Group data summarized the performance index measured with different ISI indicated at the X-axis (middle). Average performance index against the ISI, which is fitted with a sigmoid function. The coincidence time window is defined as the t50 of the sigmoidal function, and indicated with the shaded area. The dashed vertical lines at 16.6 s represents the coincidence time window of the WT flies. In (**D**), Trh mutant flies were used. In (**E**), flies were pretreated with the SSRI fluoxetine before experiment.

### 5-HT modulates the coincidence time window of circuit plasticity

A potential mechanism underlying this bi-directional behavioral modulation is that 5-HT could regulate the change of synaptic plasticity induced by odorant-shock pairing. Previous electrophysiological results suggest that pairing an odorant with dopaminergic reinforcement induces synaptic depression between KCs and the MBON-γ1pedc (Hige et al., 2015). Similar depression was observed using Ca^2+^ imaging in the MBON-γ1pedc after odorant-shock pairing (Felsenberg et al., 2018; Perisse et al., 2016). Therefore, to measure the change in plasticity before and after odorant-shock pairing in live flies, we expressed GCaMP6s in the postsynaptic MBON- γ1pedc neurons (Figure S1A). During the pairing session, a paired odorant (CS+) and electric shocks were delivered to the head-fixed fly with a 10-s ISI. Another odorant (CS-) was delivered as an unpaired stimulus (Figure S1B). In the postsynaptic MBON-γ1pedc, odorant-shock pairing significantly depressed the Ca^2+^ responses to the CS+, while the Ca^2+^ responses to the CS- remained (Figure S1C), which is consistent with previous reports (Hige et al., 2015). Given that KCs release the excitatory neurotransmitter ACh (Barnstedt et al., 2016), we then examined ACh dynamics in the γ1 compartment by expressing ACh3.0 in KCs (Figure 2A). Similar to the phenomenon observed for the postsynaptic Ca^2+^ signal, we found that odorant-shock pairing specifically reduced ACh release in response to the CS+, but had no significant effect on the CS- (Figures 2B and 2C). These findings revealed that odorant-shock pairing depresses presynaptic ACh release and the postsynaptic Ca^2+^ signal.

**Figure 2.**
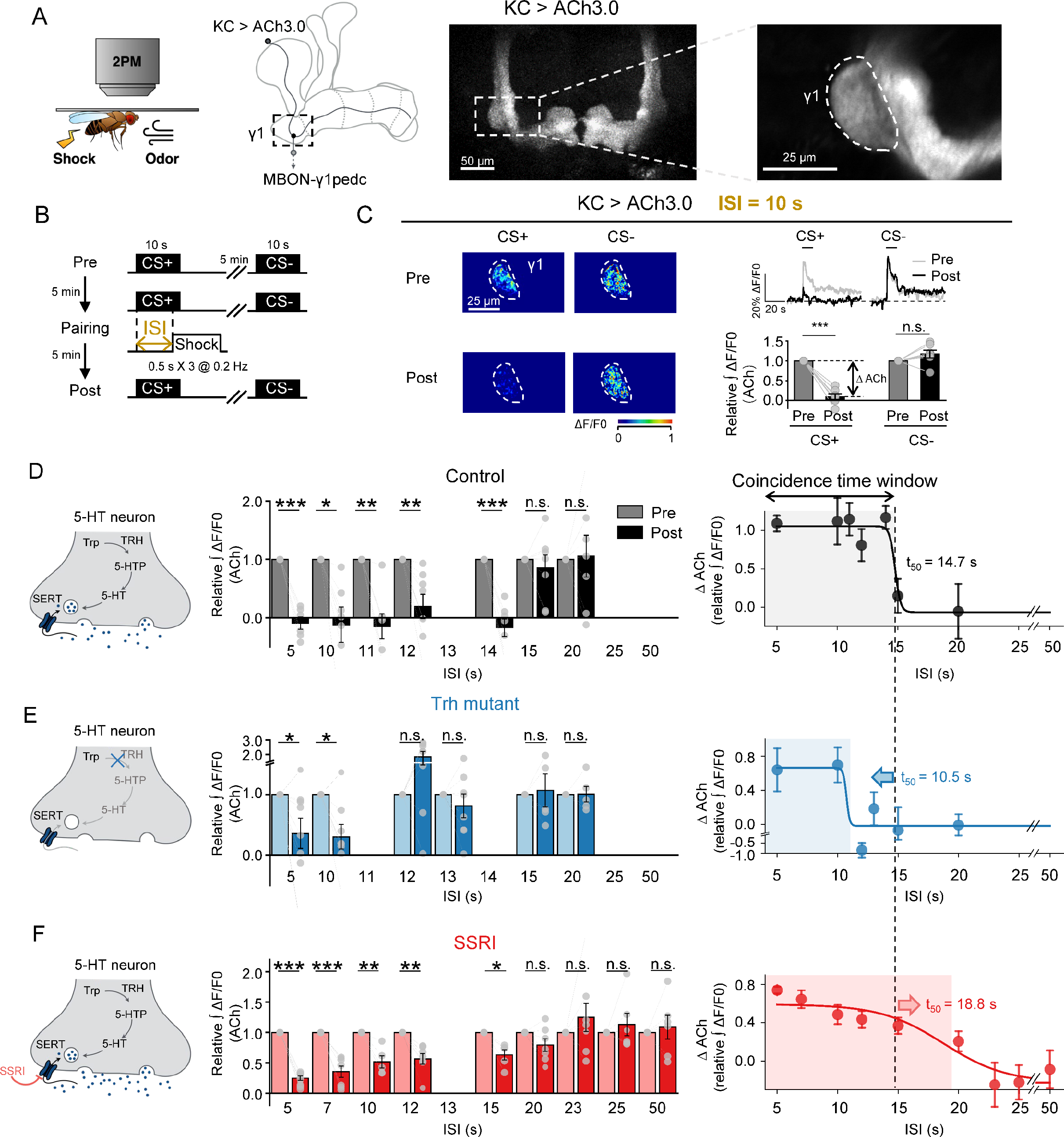
5-HT signaling can bi-directionally modulate the coincidence time window for synaptic plasticity change. (A) Schematic diagram (left and middle) depicting the strategy for measuring the synaptic plasticity changes in the γ1 compartment. ACh was measured using ACh3.0 expressed in KCs (right). (B) Schematic diagram showing the experimental protocol. (C) Representative pseudocolor images (left), average traces (top right), and group data (bottom right) showing the change in ACh3.0 fluorescence in response to the paired conditioned stimulus (CS+) and the unpaired conditioned stimulus (CS-) pre and post CS-US pairing with a 10-s ISI in control flies. (**D**-**F**) Left: schematic diagrams showing the strategy for each experiment. Middle: group relative change in ACh3.0 fluorescence in response to CS+ measured before (light) and after (dark) CS-US pairing using the indicated ISI (X-axis). Right: plot depicting the relative responses against ISI, where the ACh decrease level (ΔACh) after pairing are fitted by a sigmoid function. The coincident time window is defined as the t50 of the sigmoidal function, and indicated with the shaded area. The dashed vertical line at 14.7 s represents the coincidence time window in control flies. In (**E**), Trh mutant flies were used. In (**F**), flies were pretreated with the SSRI fluoxetine before experiment. \**p*<0.05, \*\**p*<0.01, and \*\*\**p*<0.001 (Student’s *t*-test).

To explore whether the induction of presynaptic ACh signal depression also relies on a specific coincidence time window, we systematically profiled the relationship between the ISI and synaptic plasticity change. In control flies, we found that the synaptic depression occurred only when the odorant and shock were delivered ≤14 s (Figure 2D). The t50 of the sigmoid function-fitted curve of the ACh change (Δ ACh) is 14.7 s, which is close to the 16.9-s coincidence time window for aversive learning behavior (Figure 1C). To examine whether 5-HT also regulates the coincidence time window for synaptic depression in the γ1 compartment, we profiled the time window of Trh mutant and SSRI fed flies. Consistent with our behavior results, we found that the coincidence time window in Trh mutant flies was shortened to 10.5 s (Figure 2E), while SSRI feeding slightly prolonged the coincidence time window to 18.8 s (Figure 2F). These results indicated that modulating the 5-HT level could bi-directionally regulate coincidence time windows of synaptic plasticity in the γ1 compartment of the MB.

### 5-HT signal in MB is from the DPM neuron

Each hemisphere of the *Drosophila* brain contains only one DPM neuron that innervates all three MB lobes and the peduncle region (the joint between dendrites and axons of KCs) (Figures 3A and S2). Previous studies used the Ca^2+^ indicator GCaMP or the pHluorin-based pH reporter synapto- pHluorin to indirectly measure neurotransmission from the DPM neuron, which only reflects the neuronal activity but does not dissect the role of specific neurotransmitter (Yu et al., 2005). To directly measure 5-HT release selectively from the DPM neuron, we performed *in vivo* two-photon imaging on flies expressing the green fluorescent 5-HT1.0 sensor in the KCs and the opsin CsChrimson in the DPM neuron (Figures 3A and 3B). Optogenetic stimulation induced transient changes in 5-HT1.0 fluorescence in the peduncle region and all γ lobe compartments (Figure 3C- 3G). Taking the γ2-5 compartments as examples, we found that the 5-HT1.0 response increased incrementally with light pulse number, with no notable difference among the four compartments, suggesting homogenous release ability of 5-HT at the DPM neuron’s terminals throughout these regions.

**Figure 3.**
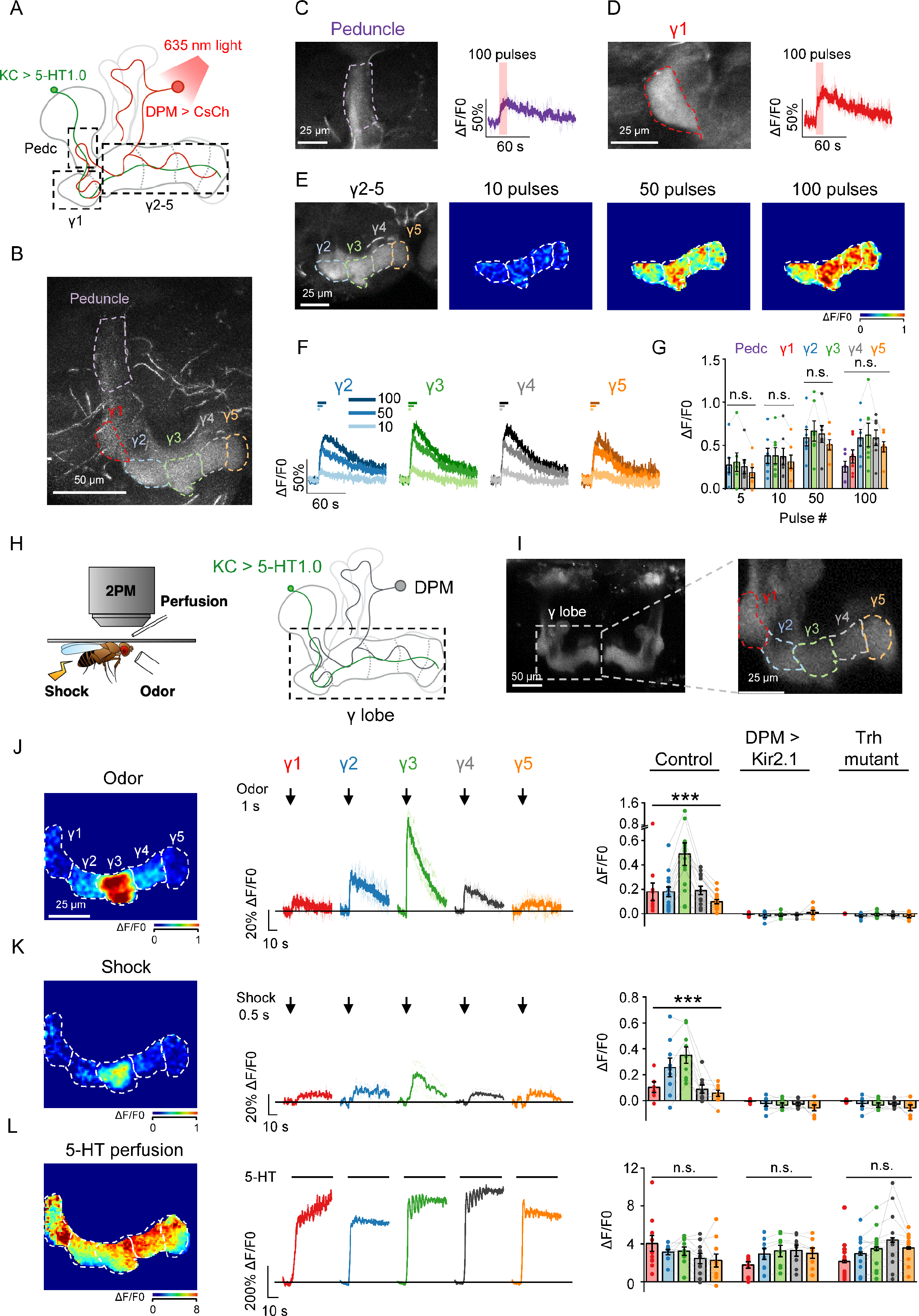
5-HT1.0 can be used to detect 5-HT release from the DPM neuron induced with optogenetics, odorant, and shock stimuli. (A) Schematic diagram depicting the experimental setup combining *in vivo* imaging with optogenetic stimulation. The CsChrimson-expressing DPM neuron (red) was activated with 1- ms pulses of 635-nm light delivered at 10 Hz, and 5-HT was measured using 5-HT1.0 expressed in KCs (green). The MB (solid line) and compartments (dashed line) of the γ lobe are shown in gray. The nicotinic ACh receptor antagonist mecamylamine (Meca, 100 μM) was present during the optogenetic experiments to avoid interference from indirect activation. (B) Representative *in vivo* fluorescence image of 5-HT1.0 expressed in KCs. (**C** and **D**) Representative fluorescence images and traces of 5-HT1.0 in the peduncle (**C**) and the γ1 compartment (**D**); where indicated, 100 light pulses were applied. (**E**-**G**) Representative fluorescence image (**E**, left panel), pseudocolor images (**E**, right panels), traces (**F**), and group data (**G**) of the change in 5-HT1.0 fluorescence in response to the indicated number of optogenetic stimuli in the different γ lobe compartments. (H) Schematic diagram depicting the experimental setup combining *in vivo* imaging with physiological stimuli and perfusion. 5-HT was measured in the γ lobe using 5-HT1.0 expressed in KCs. (I) Representative fluorescence images of 5-HT1.0 expressed in KCs. (**J**-**L**) Representative pseudocolor images (left), traces (middle), and group data (right) of the change in 5-HT1.0 fluorescence in response to a 1-s odorant (**J**), a 0.5-s electric shock (**K**), or application of 100 μM 5-HT (**L**) in control flies, flies overexpressing Kir2.1 to silence the DPM neuron, and Trh mutant flies to reduce 5-HT production. In this and subsequent figures, traces are shown as the average response (bold) with corresponding individual responses (light) measured in a single fly. In this figure, group data are presented as the mean ± SEM, overlaid with the data obtained from each fly. \**p*<0.05, \*\*\**p*<0.001, and n.s., not significant (one-way ANOVA).

Next, we used 5-HT1.0 to probe 5-HT dynamics evoked by either odorant application or electric shock (Figures 3H and 3I). We found that both odorant application (Figure 3J) and electric shock (Figure 3K) induced time-locked increases of 5-HT1.0 fluorescence in the γ lobe. Interestingly, we found that these stimuli induced responses differed among different compartments in the γ lobe of control flies, with the strongest response occurring in the γ3 compartment (Figures 3J and 3K). In contrast, optogenetic stimulation produced a relatively uniform response throughout the γ lobe (Figures 3E-3G). For Trh mutant flies, the fluorescence response was eliminated under odorant and shock stimulus, similar results were obtained when the DPM neuron was silenced by expressing the inward rectifying potassium channel Kir2.1, while direct application of 5-HT still elicited a robust response (Figure 3L). These results together demonstrate the chemical specificity of fluorescence responses and suggest that the endogenous 5-HT signal measured in MB γ lobe arises from the DPM neuron.

### The DPM neuron and KCs are reciprocally connected and functionally correlated

To better understand the 5-HT modulation on coincidence time window in MB, we explored upstream and downstream connections of DPMs. Previously, the DPM neuron was suggested to form a recurrent loop with KCs in the α’/β’ lobe (Krashes et al., 2007). However, that has not been verified experimentally. An analysis of recently published EM connectomics (Li et al., 2020; Scheffer et al., 2020) revealed that the DPM neuron forms reciprocal connections with KCs, as well as other cell types, including DANs in the paired posterior lateral 1 (PPL1) cluster, DANs in the protocerebral anterior medial (PAM) cluster and a single GABAergic anterior paired lateral (APL) neuron (Figures S3A, S3B, S3D, and S3E). Furthermore, both the input and output synapses of the DPM neuron are distributed in all compartments of the MB. By analyzing the percentile from each cell type, we found that more than 80% of the DPM’s upstream cells are KCs and KCs comprise more than 50% of the DPM’s downstream cells (Figures S3B and S3E). Moreover, we found that all 1931 KCs examined in our analysis form reciprocal connections with the DPM neuron. On average, each KC has 28 pre-synapses and 16 post-synapses that are connected with the DPM neuron (Figures S3, S3C, S3F and S3G).

To further examine the functional relationship between the DPM and KCs (Figure S4A), we used ACh3.0 to measure ACh release from KCs. Additionally, we used GCaMP5 and 5-HT1.0 to measure the DPM neuronal activity and 5-HT release from the DPM neuron. We performed *in vivo* two-photon imaging in the γ2-5 compartments in flies expressing each sensor, while applying an odorant or electric shock stimuli. By comparing the resulting patterns, we found that ACh dynamics are positively correlated with the Ca^2+^ signal in the DPM neuron and 5-HT dynamics (Figures S4B and S4C), suggesting that the DPM neuron and KCs are both reciprocally connected and functionally correlated.

### KCs are both necessary and sufficient for activating the DPM neuron

To figure out the input-output relationship between the DPM and KCs, we generated transgenic flies expressing both the inhibitory DREADD (Designer Receptor Exclusively Activated by Designer Drugs) hM4Di (Armbruster et al., 2007; Becnel et al., 2013; Roth, 2016) and 5-HT1.0 in KCs (Figure 4A). When the hM4Di agonist deschloroclozapine (DCZ) (Nagai et al., 2020) was applied to suppress KCs activity, we found that the odor- and shock-induced 5-HT release in the γ lobe was abolished (Figures 4B and 4C), suggesting that KC excitatory input is required for the 5-HT release from the DPM neuron during odor and shock stimulations.

**Figure 4.**
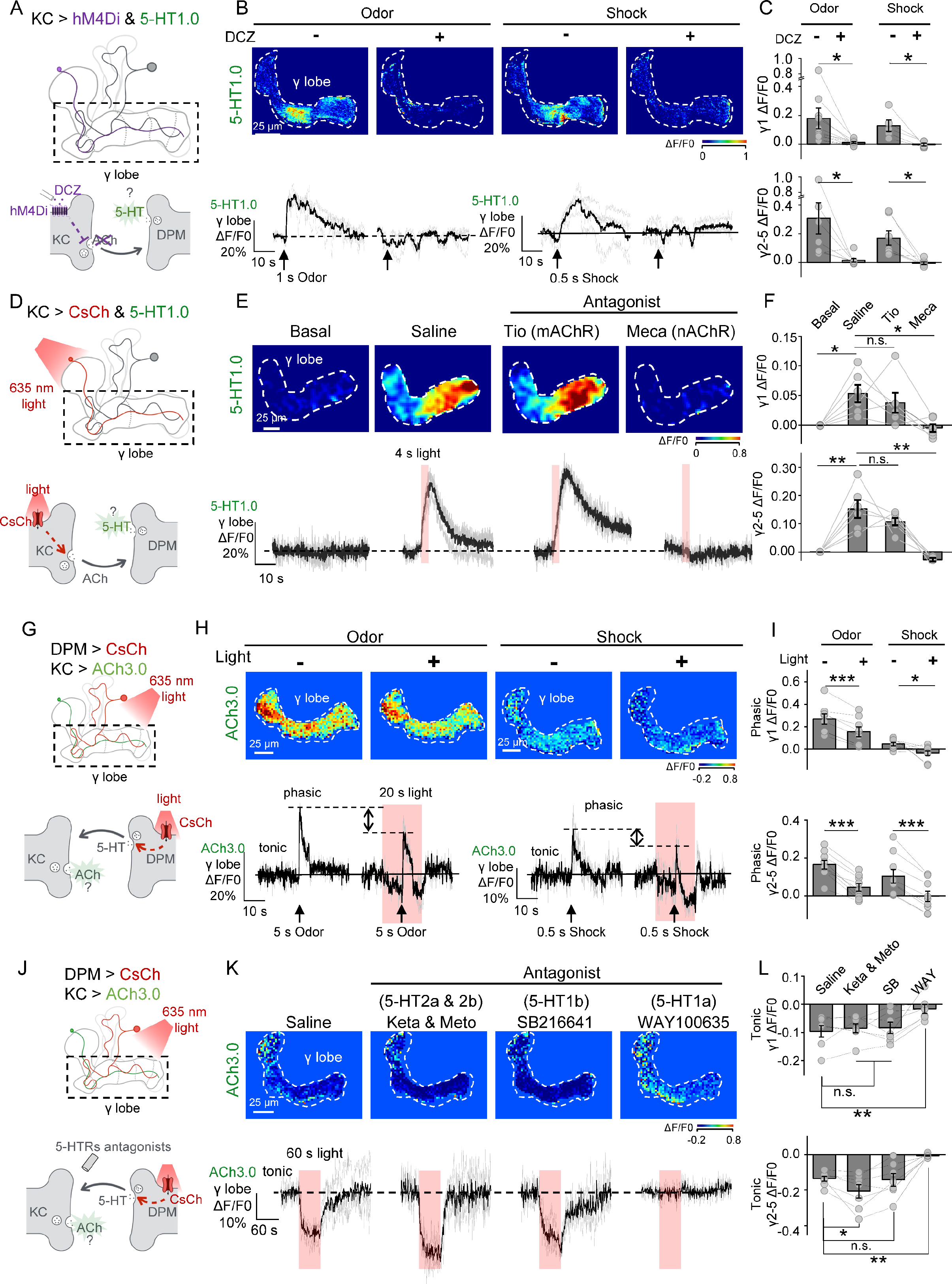
5-HT release from the DPM neuron is induced by ACh release from KCs and provides inhibitory feedback to KCs. (**A**) Schematic diagram depicting the setup used for the experiments shown in (**B**) and (**C**). hM4Di-expressing KCs were silenced by applying 30 nM deschloroclozapine (DCZ), and 5-HT was measured in the γ lobe using 5-HT1.0 expressed in KCs. (**B** and **C**) Representative pseudocolor images (**B**, top), traces (**B**, bottom), and group data (**C**) of the change in 5-HT1.0 fluorescence in response to a 1-s odorant application or 0.5-s electric shock in the absence or presence of 30 nM DCZ. (**D**) Schematic diagram depicting the setup used for the subsequent experiments. CsChrimson- expressing KCs were activated by 40 1-ms pulses of 635-nm light applied at 10 Hz, and 5-HT was measured in the γ lobe using 5-HT1.0 expressed in KCs. (**E** and **F**) Representative pseudocolor images (**E**, top), traces (**E**, bottom), and group data (**F**) of the change in 5-HT1.0 fluorescence in response to optogenetic stimulation in saline, the muscarinic ACh receptor antagonist Tio (100 μM), or the nicotinic ACh receptor antagonist Meca (100 μM). (**G**) Schematic diagram depicting the experimental setup for the subsequent experiments. The CsChrimson-expressing DPM neuron was activated using 1-ms pulses of 635-nm light at 10 Hz, and ACh was measured in the γ lobe using ACh3.0 expressed in KCs. (**H** and **I**) Representative pseudocolor images (**H**, top), traces (**H**, bottom), and group data (**I**) of the change in ACh3.0 fluorescence in response to a 1-s odorant application or 0.5-s electric shock either with or without a 20-s optogenetic stimulation. (**G**) Schematic diagram depicting the experimental setup for the subsequent experiments. Similar to (**J**), but different 5-HT receptor antagonists are applied. (**K** and **L**) Representative pseudocolor images (**K**, top), traces (**K**, bottom), and group data (**L**) of the change in ACh3.0 fluorescence in response to a 60-s optogenetic stimulation. Different compounds were sequentially added into the bath solution without washing, including the 5- HT2a antagonist ketanserin (Keta), the 5-HT2b antagonist metoclopramide (Meto), the 5-HT1b antagonist SB216641, and the 5-HT1a antagonist WAY100635 (all applied at 20 μM each). In this figure, group data are presented as the mean ± SEM, overlaid with the data obtained from each fly. \**p*<0.05, \*\**p*<0.01, \*\*\**p*<0.001, and n.s., not significant (Student’s *t*-test). For these experiments in (**D** - **L**), the gap junction blocker CBX (100 μM) was included.

Next, we examined whether ACh is sufficient to activate the DPM neuron (Figure S5A). We found that perfusing ACh on the horizontal lobe induced an increase in 5-HT1.0 fluorescence that can be blocked by the nicotinic ACh receptor (nAChR) antagonist mecamylamine (Meca) (Figures S5B and S5C), which is consistent with recent transcriptomics data showing that nicotinic ACh receptors, but not muscarinic receptors (mAChR), are expressed in the DPM neuron (Figure S6A (Aso et al., 2019)). Importantly, adding other neurotransmitters such as DA, OA, glutamate (Glu) or GABA in the presence of Meca also did not cause an increase in 5-HT1.0 fluorescence, whereas application of 5-HT elicited a robust response (Figures S5B and S5C). Thus, ACh provides the excitatory input to the DPM neuron.

Because externally ACh perfusion lacks cell type specificity, we further examined whether selectively activating KCs is sufficient to trigger the release of 5-HT from the DPM neuron. We therefore expressed CsChrimson and 5-HT1.0 in KCs (Figure 4D). Optogenetic activation of KCs induced a 5-HT signal in the γ lobe (Figures 4E, 4F and S7) and this signal can be blocked by the nAChR antagonist Meca but not the mAChR antagonist tiotropium (Tio). In addition, we used a 2- photon laser to activate a specific region of the MB and observed localized 5-HT release (Figure S8). These results indicate that activation of KCs is both necessary and sufficient to drive the localized release of 5-HT from the DPM neuron, and this effect is mediated by nAChRs.

### The DPM neuron provides inhibitory feedback to the KCs

Besides the KCs to the DPM neuron regulation, we next examined the effect of 5-HT released from the DPM neuron on KCs. We therefore expressed the CsChrimson to optogenetically activate the DPM neuron, with ACh3.0 in the KCs to measure both basal and stimuli-evoked fluorescent signals, indicating tonic and phasic ACh dynamics respectively (Figure 4G). Because the DPM neuron is connected to a GABAergic APL neuron via gap junctions, we used the gap junction blocker carbenoxolone (CBX) to prevent indirect activation of the APL neuron (Connors, 2012). In the absence of optogenetic stimulation, application of either odorant or electric shock induced phasic ACh release in the γ lobe, and these responses were significantly reduced when the stimuli (i.e. odor or shock) were presented 10 s after shinning the red light (Figures 4H and 4I). This DPM- activation evoked inhibitory effect was largely abolished in Trh mutant flies (Figure S9A-9C). Moreover, both the odor and shock evoked ACh release in MB were significantly increased in Trh mutant flies (Figure S9D and S9E). These two lines of evidences strengthen the inhibitory tone of 5-HT in the MB.

It has been documented that KCs show abundant neuronal activity in the absence of odor stimulation(Turner et al., 2008). Therefore, we measured the tonic ACh signal, and found it was reduced by activation of the DPM neuron (Figures 4H and 4I). 5-HT mediated inhibition to ACh release was largely abolished in Trh mutant flies. Analysis of recent transcriptomic data (Aso et al., 2019) revealed that both the 5-HT1a and 5-HT1b receptors are expressed in KCs in the γ lobe (Figure S6B). Both receptor subtypes are coupled to the inhibitory Gαi pathway (Saudou et al., 1992). Therefore, to determine which 5-HT receptor subtype mediated inhibitory 5-HT signaling to KCs, we applied 5-HT receptor subtype specific antagonists (Suzuki et al., 2020) and found that blocking the 5-HT1a receptor with WAY100635 prevented the optogenetically induced decrease of tonic ACh signaling. In contrast, blocking the 5-HT1b, 5-HT2a, or 5-HT2b receptor had no such effects (Figures 4J-4L). Taken together, these functional results reveal a reciprocal relationship between the DPM neuron and KCs in the γ lobe, in which KCs release ACh to locally activate the DPM neurons, while the DPM neuron releases 5-HT to inhibit ACh release via the 5-HT1a receptor.

### DPM-mediated serotonergic feedback inhibition modulates the coincidence time window

Having established functional relationships between the DPM neuron and KCs, we then examined the role of serotonergic inhibitory feedback for synaptic plasticity change in the γ1 compartment revealed by ACh3.0 imaging (Figures 5A and 5B). By specifically silencing the DPM neuron with Kir2.1, we found that the coincidence time window was shortened to 10.9 s (Figures 5C, 2E). Whereas the optogenetical activation of the DPM neuron with CsChrimson significantly prolonged the coincidence time window to 24.0 s (Figure 5D). To demonstrate the necessity of 5-HT metabolism specifically in the DPM neuron, we conducted optogenetic stimulation with Trh mutant flies and yielded an 11.2-s coincidence time window, which was similar to that found in Trh mutant and DPM silenced flies (Figure 5E). Moreover, the coincidence time windows were shortened when we mutated the 5-HT1a receptor (Qian et al., 2017) (Figure 5F) or knocked down its expression in KCs with RNAi (Figure 5G) (12.3 s for 5-HT1a mutant flies, and 12.2 s for 5-HT1a RNAi flies respectively).

**Figure 5.**
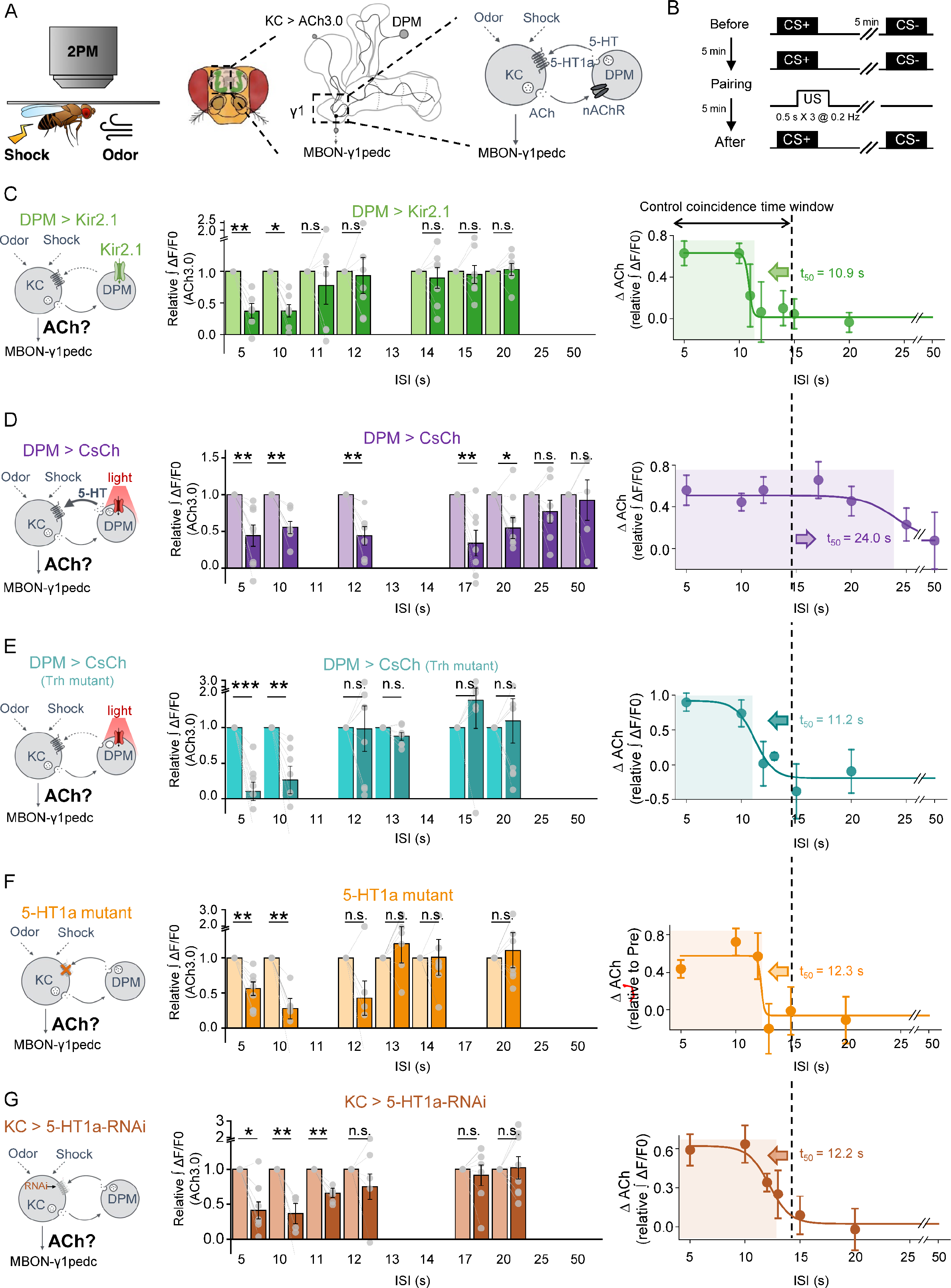
5-HT signals from DPM can bi-directionally modulate the coincidence time window for changing synaptic plasticity. (A) Schematic diagram (left and middle) depicting the strategy for measuring the effect of DPM-mediated serotonergic inhibitory feedback on changes in synaptic plasticity in the γ1 compartment. ACh was measured using ACh3.0 expressed in KCs (right). (B) Schematic diagram showing the experimental protocol. (**C**-**G**) Left: schematic diagrams showing the strategy for each experiment. Middle: group relative change in ACh3.0 fluorescence in response to CS+ measured before (light) and after (dark) CS-US pairing using the indicated ISI. Right: plot depicting the relative depression of ACh signals in response to CS+ against ISI, where the decreases are fitted by a sigmoid function. The coincident time window is defined as the t_50_ of the sigmoidal function, and indicated with the shaded area. The dashed vertical line at 14.7 s represents the coincidence time window in control flies. In (**C**), the DPM neuron expressed Kir2.1. In (**D**), the DPM neuron expressed CsChrimson, which was activated using 10-ms pulses of 635-nm light at 4 Hz, applied from the start of odorant application to 4.5 s after electric shocks were applied. In (**E**), the DPM expressed CsChrimson in Trh mutant flies, which was activated using identical protocols as in (**D**). In (**F**), the 5-HT1a receptor was mutated. In (**G**), the 5-HT1a receptor was knocked down in KCs with RNAi. Data fitted with a nonlinear Dose-Response function. In this figure, group data are presented as the mean ± SEM, overlaid with the data obtained from each fly. \**p*<0.05, \*\**p*<0.01, and \*\*\**p*<0.001 (Student’s *t*-test).

Finally, we wanted to confirm whether the time regulating function of DPM-mediated serotonergic feedback inhibition holds true for the learning process. (Figures 6, A and B). For DPM neuron silenced flies, the coincidence time window was shortened to 10.5 s (Figure 6C). Whereas the time window was prolonged to 44.1 s for the DPM neuron activated group (Figure 6D). When we specifically expressed the TRH in the DPM neuron of Trh mutant flies, interestingly, we found the coincidence time window was not only rescued but further prolonged to 33.4 s, supporting the sufficiency of 5-HT signal from the DPM neuron (Figure 6E). Systematically Mutating 5-HT1a or specifically knocking down the 5-HT1a in KCs shortened the coincidence time window to 14.7 s and 10.6 s respectively (Figures 6F and 6G).

**Figure 6.**
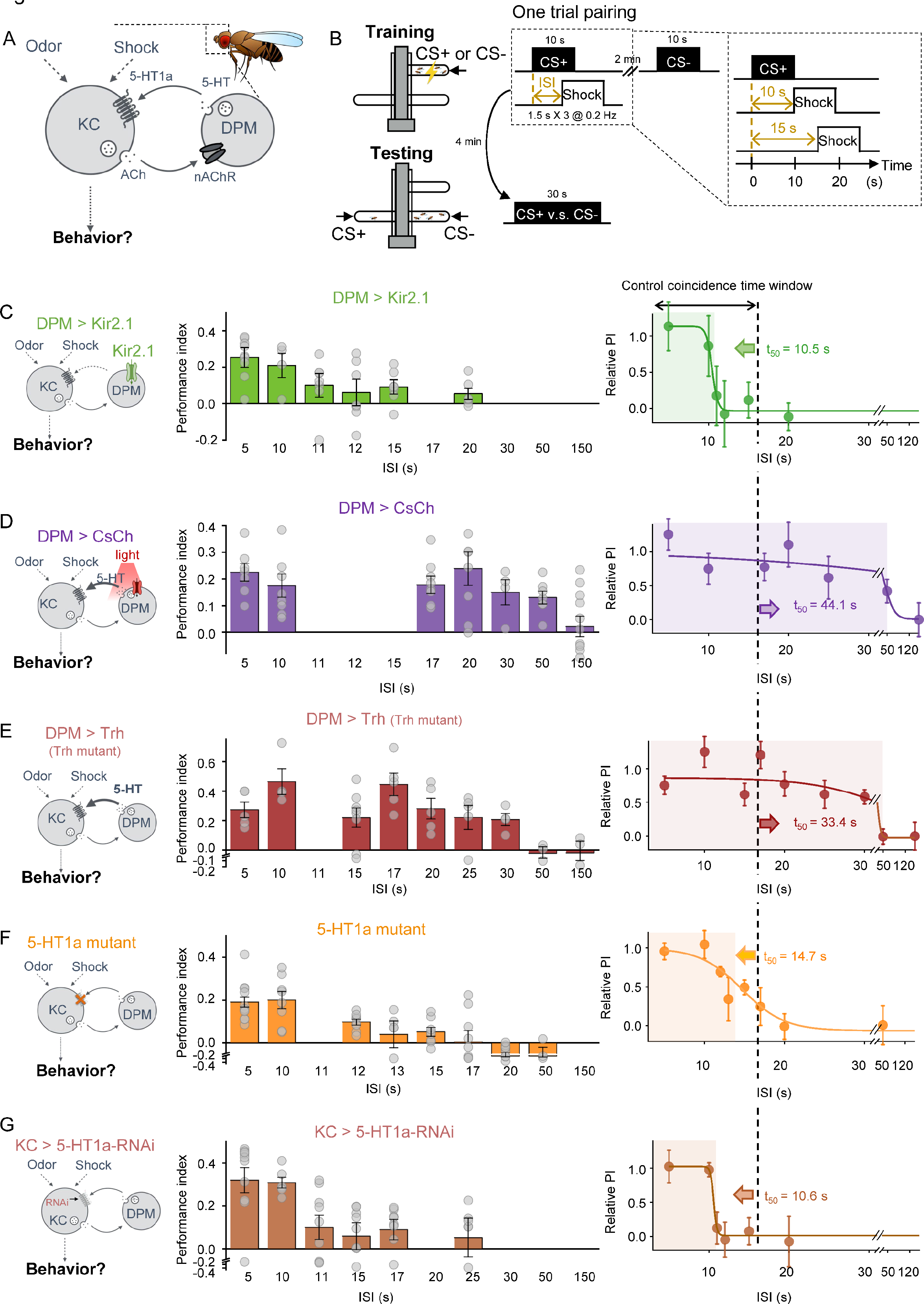
5-HT signaling can bi-directionally modulate the coincidence time window of olfactory learning. (A) Schematic diagram depicting the DPM-mediated inhibitory serotonergic feedback to KCs. (B) T-maze protocol for measuring how the inter-stimulus interval (ISI) affects odorant-shock pairing-induced aversive memory. (**C**-**G**) Left: schematic diagrams showing the strategy for each experiment. Middle: group data summarizing the performance index measured using the indicated ISI. Right: plot depicting the averaged relative performance index against the ISI, which is fitted with a sigmoid function. The coincident time window is defined as the t_50_ of the sigmoidal function, and indicated with the shaded area. The dashed vertical line at 16.5 s represents the coincidence time window of the control flies. In (**C**), the DPM neuron expressed Kir2.1. In (**D**), the DPM neuron expressed CsChrimsn, which was activated with continuous 635-nm light applied from the beginning of the odorant application to 3.5 s after the electric shocks were applied. In (**E**), the Trh was conditional over-expressed in DPM in Trh mutant flies. In (**F**), the 5-HT1a receptor was mutated. In (**G**), the 5-HT1a receptor was knocked down in KCs with RNAi. Data in **C**-**G** are fitted with a nonlinear Dose-Response function.

Taken together, our results indicate that modulating the DPM activity or 5-HT signal yields shifted coincidence time windows of synaptic plasticity in the γ1 compartment of the MB, which are positively correlated with the coincidence time windows of the learning behavior (Figure 7A). Meanwhile, the learning ability as well as the amplitude of the ACh depression is not affected (Figure 7B). In summary, the 5-HT signal from the DPM neuron selectively serves as a specific timing modulator to regulate the coincidence time window in the olfactory associative learning process (Figure 7C).

**Figure 7.**
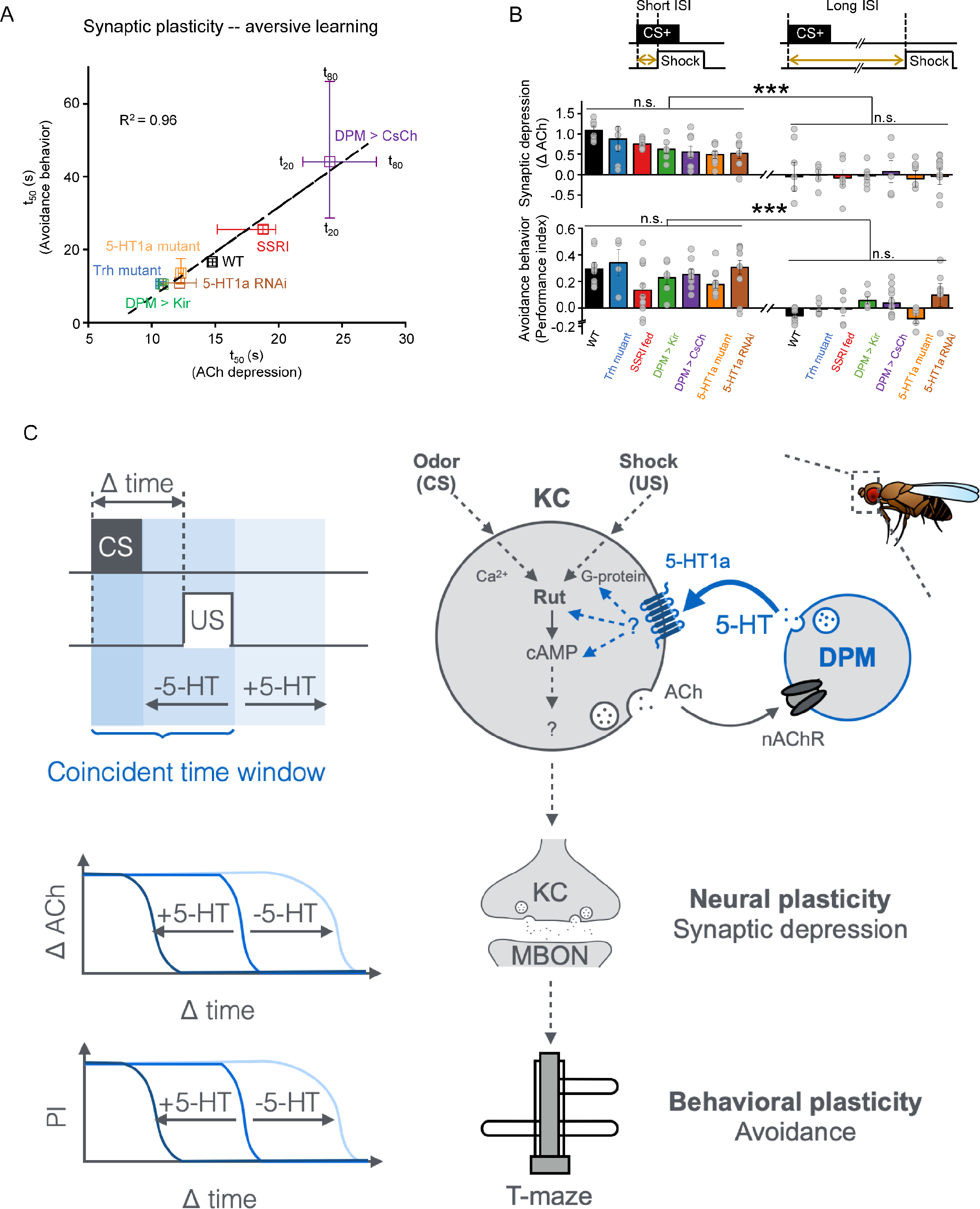
5-HT signal bi-directionally regulates the coincidence time window of associative learning. (A) Correlation analysis of coincidence time windows (t50) between synaptic plasticity (X-axis) and aversive learning performance (Y-axis) and synaptic plasticity of indicated fly groups. Error bars indicate the temporal range from t20 to t80. The data were fit to a linear function, with the corresponding correlation coefficients shown. (B) Comparing the amplitudes of behavioral avoidance and synaptic depression with different temporal range of indicated fly groups. Short ISI: data of avoidance behavior and synaptic plasticity are quantified when ISI = 5 s for all fly groups. Long ISI: data of avoidance behavior are quantified when ISI = 20 s for WT, Trh mutant and DPM > Kir2.1, and when ISI = 150 s for DPM > CsChrimson and SSRI; data of synaptic plasticity are quantified when ISI = 20 s for WT, Trh01 and DPM > Kir2.1 when ISI = 50 s for DPM > CsChrimson and when ISI = 50 for SSRI. (**C**)Working model depicting the mechanism by which local 5-HT signaling can bi-directionally modulate the coincidence time window of associative learning. In the *Drosophila* olfactory associative learning center, the Kenyon cells (KCs) receive inhibitory feedback from a single serotonergic dorsal paired medial (DPM) neuron. The KC innervates the mushroom body output neurons (MBONs). Pairing between the conditioned stimulus (CS) and the unconditioned stimulus (CS) regulating the coincidence time window for the change in synaptic plasticity and subsequent learning behavior. Data in **A** and **B** are re-organized from Fig. 1, 2, 5 and 6. Data presented in **B** as the mean ± SEM. n.s., no significant difference. ***., p < 0.001 (One-way ANOVA).

## Discussion

Nearly a century ago, Ivan Pavlov proposed the associative conditioning theory, stating that “A … most essential requisite for … a new conditioned reflex lies in a coincidence in time of … the neutral stimulus with … unconditioned stimulus” (Pavlov and Anrep, 1927). Here, we reported that the coincidence time window between CS and US for olfactory learning of Drosophila could be bi- directionally regulated by 5-HT signal. We further dissected the microcircuit in the MB, where the DPM neuron releases 5-HT to provide inhibitory feedback to KCs. These results support a circuitry model in which the animal can maintain a physiologically precise time window to extract meaningful associations from the surrounding environment.

### Serotonergic neuromodulation in the olfactory mushroom body

Despite the known importance of serotonergic signaling in olfactory learning in *Drosophila* (Ganguly et al., 2020; Johnson et al., 2011; Keene et al., 2006; Keene et al., 2004; Krashes et al., 2007; Lee et al., 2011; Sitaraman et al., 2008; Waddell et al., 2000; Wu et al., 2011; Yu et al., 2005), the dynamics of 5-HT signaling *in vivo* and the mechanisms that regulate this signaling processes are largely unknown. Previously, addressing these fundamental biological questions has been difficult due to the absence of suitable tools for monitoring 5-HT dynamics *in vivo* with high spatiotemporal resolution. Using our 5-HT1.0 sensor, we measured 5-HT release in specific compartments in the MB γ lobe in response to odor application (CS) and electric shock (US), which is regulated by local ACh release from KCs. Each hemisphere contains at least three serotonergic neurons that project to the MB, the DPM neuron innervates all lobes and the peduncle, the serotonergic projection neuron (SPN) innervates only the peduncle (Scheunemann et al., 2018), and the contralaterally-projecting serotonin-immunoreactive deuterocerebral interneuron (CSDn) innervates the calyx (Coates et al., 2020; Coates et al., 2017; Dacks et al., 2006; Suzuki et al., 2020; Zhang et al., 2019a). However, our finding that the physiological stimulation-evoked increase in 5-HT1.0 fluorescence in the γ lobe disappeared when the DPM neuron was silenced suggests that the DPM neuron is the principal source of 5-HT release in the γ lobe.

### Inhibitory feedback circuits in the learning center

Based on previous light microscopy images and behavioral studies, the DPM neuron and KCs are believed to form a recurrent loop in the α’/β’ lobe (Krashes et al., 2007), and this notion is supported by EM connectomics (Li et al., 2020; Scheffer et al., 2020). In addition to this structural connection, our functional imaging results reveal that the DPM neuron provides inhibitory feedback to KCs. Although the DPM neuron has been shown to release both 5-HT and GABA (Haynes et al., 2015), our results indicate that the inhibitory effect on KCs, which regulates the coincidence time window, is mediated primarily by 5-HT acting on 5-HT1a receptors in the KCs.

Each hemisphere contains a GABAergic APL neuron with neuropils that ramify throughout the MB, including the calyx (Liu and Davis, 2009). The APL is not only anatomically similar to the DPM neuron, but functionally the APL also forms reciprocal connections with KCs and provides inhibitory feedback (Amin et al., 2020; Inada et al., 2017; Papadopoulou et al., 2011; Wu et al., 2012). Moreover, GABAA receptors-mediated inhibitory feedback can control the sparseness of odorant coding in KCs, which allows the animal to discriminate between similar odors (Lei et al., 2013; Lin et al., 2014). Here, our report that the DPM-mediated serotonergic inhibitory feedback regulates the coincidence time window between stimuli. Given that 5-HT and GABA signals in MB operate in parallel to regulate the time window and sparseness of odorant coding (Lee et al., 2011) respectively, MB likely recruits two inhibitory feedback signals in order to execute orthogonal functions of learning.

### Odorant-shock pairing induces presynaptic depression

A large number of studies reported a wide range of olfactory learning‒related changes in synaptic plasticity in the *Drosophila* MB (Akalal et al., 2010; Berry et al., 2018; Bilz et al., 2020; Boto et al., 2014; Boto et al., 2019; Bouzaiane et al., 2015; Cohn et al., 2015; Dylla et al., 2017; Felsenberg et al., 2017; Felsenberg et al., 2018; Gervasi et al., 2010; Handler et al., 2019; Hige et al., 2015; Louis et al., 2018; McCurdy et al., 2021; Owald et al., 2015; Perisse et al., 2016; Placais et al., 2013; Sabandal et al., 2021; Sejourne et al., 2011; Stahl et al., 2021; Wang et al., 2008; Yu et al., 2006; Yu et al., 2005; Zhang and Roman, 2013; Zhang et al., 2019b; Zhou et al., 2019). However, some studies differed with respect to the location (i.e., the specific MB compartment), direction (i.e., potentiation vs. depression) and whether the change occurs in presynaptic KCs or postsynaptic MBONs. By performing *in vivo* imaging with ACh3.0 and GCaMP, we found that odorant-shock pairing induces depression of the ACh signal released from KCs and Ca^2+^ signal within the MBON- γ1pedc. In addition, we found that postsynaptic Ca^2+^ responses to the CS- are unaffected by odorant-shock pairing, suggesting that the change in synaptic plasticity is more likely to occur in the presynaptic KCs.

### Regulating the coincidence time window

Activities of the DPM neuron are reported to be required only for consolidating middle-term memory (i.e., 3-hour) but not for short-term memory (Keene et al., 2004; Lee et al., 2011; Yu et al., 2005). Previous studies were performed with an overlapped CS-US pairing protocol, meaning that the ISI is shorter than 10 s. This work focuses on short-term memory, and we found that the 5-HT released from the DPM neuron specifically regulated the coincidence time window. In accordance with previous studies, we found that 5-HT does not affect magnitudes of performance index and synaptic plasticity when the ISI is ≤10 s (Figure 7B). However, when the ISI >10 s, learning differences emerged between fly groups. Given that the CS was delivered for 10 s during odorant- shock pairing, it seems reasonable to speculate that the serotonergic DPM circuitry is involved primarily in trace conditioning when a temporal gap exists between the CS and US (Shuai et al., 2011). In nature, flies do not experience precisely controlled CS and US as in the lab. Their learning needs to be flexible to different CS/US regimes. Thus, the serotonin modulation extends the ability of the flies to learn in nature and improves their chance of successfully determining cause and effect.

At the neural circuit level, we found that 5-HT from the DPM neuron can bi-directionally regulate the coincidence time window of synaptic depression in the γ1 compartment, which partially explains our behavioral results. However, olfactory learning is the net result of synaptic plasticity changes in 15 MB compartments (Hige, 2018; Waddell, 2016) and each compartment has a specific set of learning rules (Aso and Rubin, 2016). Thus, whether 5-HT plays a general role in regulating timing in distinct compartments remains an open question.

Our findings prompt a series of questions about the physical basis for the coincidence time window and the role 5-HT modulation of KCs plays in extending or reducing the window. We propose two classes of hypotheses. One hypothesis is that the time window is documented by the CS-induced Ca^2+^ activity in KCs. According to previous studies, adenylyl cyclase Rutabaga detects the coincidence of odor-induced Ca^2+^ and shock-induced dopamine signal (Davis et al., 1995; Dudai et al., 1976; Dudai et al., 1985; Gervasi et al., 2010; Levin et al., 1992; Livingstone et al., 1984; Tomchik and Davis, 2009), and increases cAMP levels, therefore modulating synaptic plasticity (Figure 7C). However, we find it difficult to fit the 5-HT signal directly into this model, as activating the DPM neuron inhibits ACh release from KCs (Figure 4G-4L), and Gα/i-coupled 5-HT1a curbs the learning-related cAMP signal, both of which shorten the window. The other hypothesis is that the coincidence time window is biochemical, for example the CaMKII autophosphorylation activity, which also determines the copulation duration of *Drosophila* (Thornquist et al., 2020; Thornquist et al., 2021). It would then imply that 5-HT can somehow prolong the CaMKII autophosphorylation states. There are many interesting unknowns that can perhaps be resolved by imaging intracellular signaling cascades in KCs in the future.

In mammals, the serotonergic system plays a critical role in cognition and serves as a pharmacological target for various hallucinogens and antidepressants. A growing body of evidence suggests that 5-HT affects the perception of time and the temporal control of various behaviors (Buhot et al., 2000; Harmer et al., 2002; Meneses, 1999; Park et al., 1994; Wittmann et al., 2007). Moreover, recent rodent studies involving associative learning paradigms found that tonic 5-HT signaling encodes “patience”, as artificially inhibiting or activating serotonergic neurons can bi- directionally regulate the time that animal waits between the CS and the US (Fonseca et al., 2015; Li et al., 2016; Liu et al., 2020b; Lottem et al., 2018; Miyazaki et al., 2011a, 2012a; Miyazaki et al., 2011b, 2012b; Miyazaki et al., 2014). In our study, 5-HT also bi-directionally regulates the coincidence timing between the CS and US. In addition, studies of the rabbit nictitating membrane response found that the hallucinogen LSD (lysergic acid diethylamide, or “acid”), a non-selective 5-HT receptor agonist, can facilitate learning when the ISI is outside of the optimal range (Harvey, 2003; Harvey et al., 1988). This finding is reminiscent of our observations in *Drosophila* that the SSRI can increase learning when the ISI exceeds the optimal coincidence time window. Thus, a similar serotonergic neuromodulatory mechanism may be used in both vertebrates and invertebrates to modulate the timing of associative learning.

## Materials and Methods

### Materials

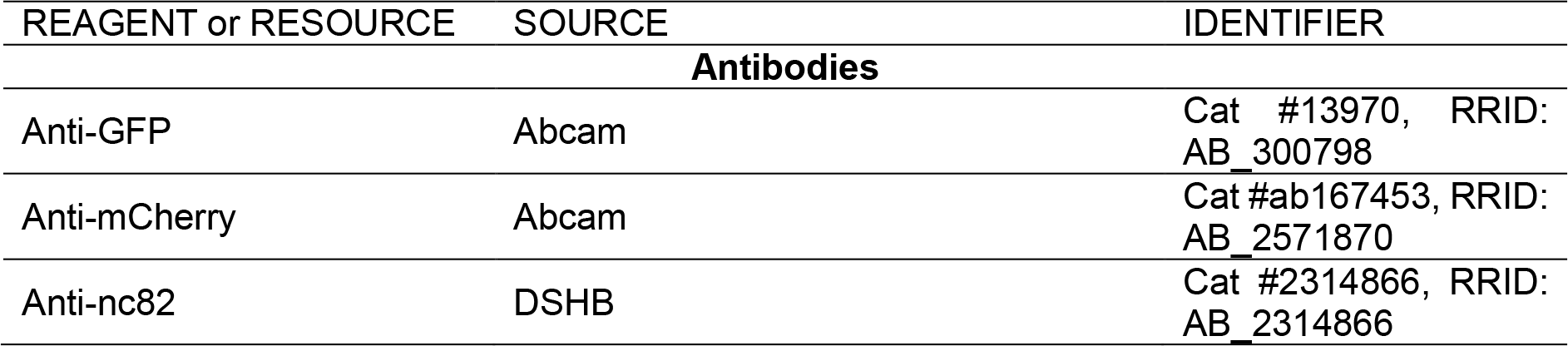
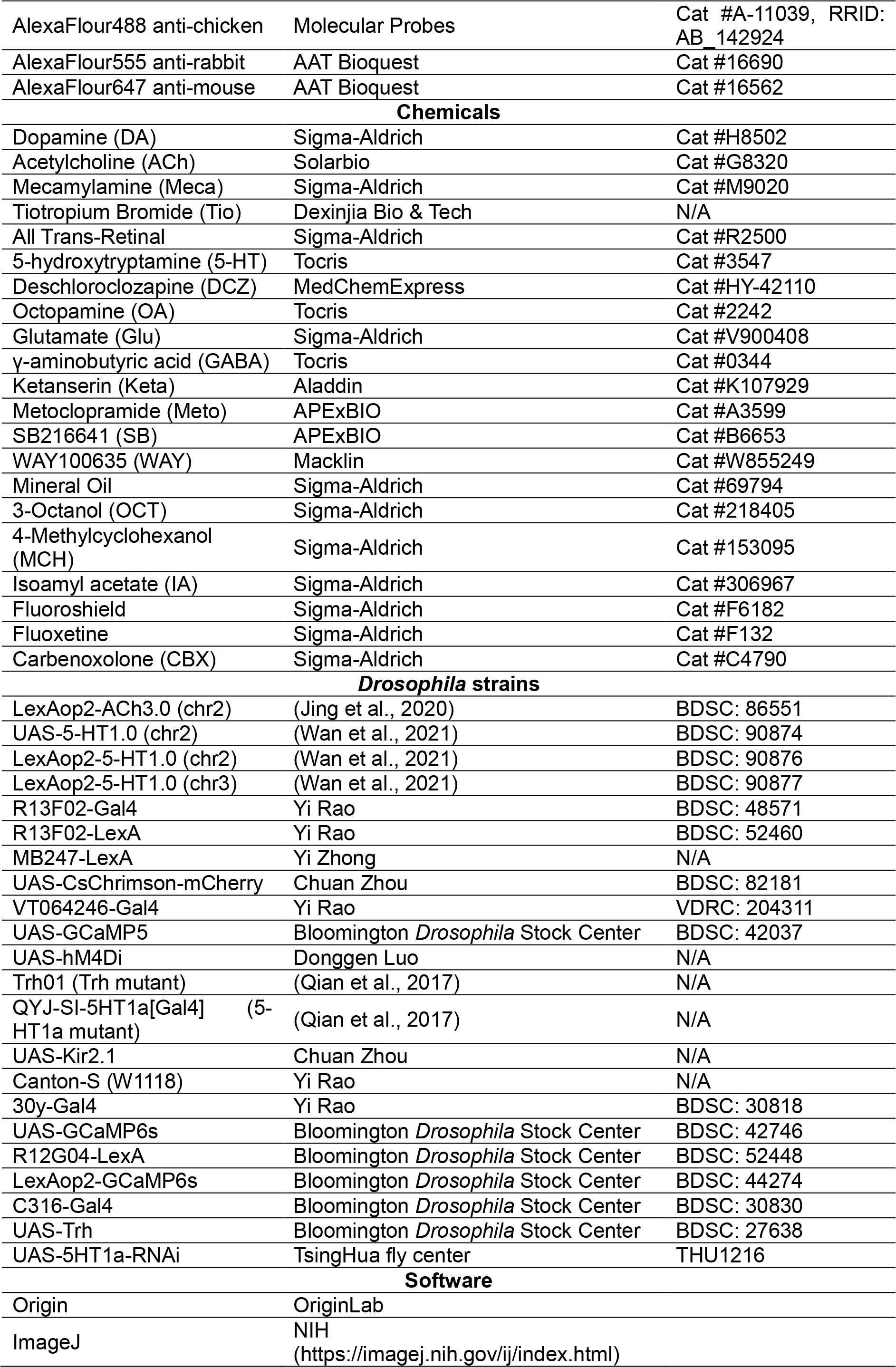
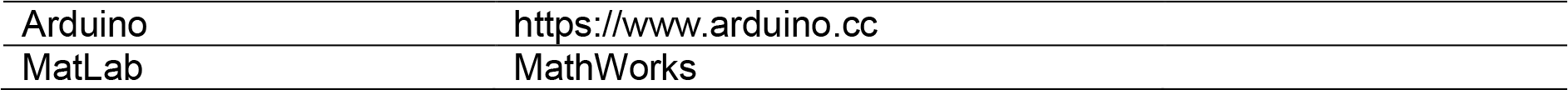

### Experiment model and subject details Flies

Transgenic flies were raised on corn meal at 25°C in 50% humidity, under a 12-hour light/12-hour dark cycle. For optogenetics, flies were transferred to corn meal containing 400 μM all-*trans*-retinal after eclosion and raised in the dark for 1-3 days before performing functional imaging and behavioral experiments. For fluoxetine feeding, flies were transferred to a tube containing a filter paper loaded with 150 μl 5% sucrose solution with 10 mM fluoxetine for 14 hours before performing behavioral experiments.

The following fly strains were used in the experiments corresponding to the following figures.

Figure 1

Canton-S (control and SSRI groups)

Trh01 / Trh01

Figure 2, Figure S1 and Figure S2

UAS-GCaMP6s / +; 30y-Gal4 / + R12G04-LexA / CyO; LexAop2-GCaMP6s / TM2 LexAop2-ACh3.0 / CyO; MB247-LexA / TM6B (control and SSRI groups) R13F02-LexA / LexAop2-ACh3.0; Trh01 / Trh01

Figure 3 and Figure S2

UAS-CsChrimson-mCherry / R13F02-LexA; VT064246-Gal4 / LexAop2-5HT1.0 UAS-5HT1.0 / CyO; R13F02-Gal4 / TM2 UAS-Kir2.1 / R13F02-LexA; VT064246-Gal4 / LexAop2-5HT1.0 R13F02-LexA / LexAop2-5HT1.0; Trh01 / Trh01

Figure 4 and Figure S4-8

LexAop2-ACh3.0 / CyO; MB247-LexA / TM6B

UAS-GCaMP5 / CyO; VT064246-Gal4 / TM6B

UAS-5HT1.0 / CyO; C316-Gal4 / TM2

UAS-hM4Di / +; UAS-5HT1.0 / +; R13F02-Gal4 / + UAS-CsChrimson-mCherry / R13F02-LexA; 30y-Gal4 / LexAop2-5HT1.0 UAS-5HT1.0 / CyO; R13F02-Gal4/TM2 LexAop2-ACh3.0 / UAS-CsChrimson-mCherry; MB247-LexA / VT064246-Gal4

LexAop2-ACh3.0 / UAS-CsChrimson-mCherry; MB247-LexA, Trh01 / VT064246-Gal4, Trh01

Figure 5

UAS-Kir2.1 / LexAop2-ACh3.0; VT064246-Gal4 / MB247-LexA

UAS-CsChrimson-mCherry / LexAop2-ACh3.0; VT064246-Gal4/ MB247-LexA

LexAop2-ACh3.0 / UAS-CsChrimson-mCherry; MB247-LexA, Trh01 / VT064246-Gal4, Trh01

LexAop-ACh3.0/+; MB247-LexA, 30y-Gal4/UAS-5-HT1a-RNAi

QYJ-SI-5HT1a[Gal4]/ QYJ-SI-5HT1a[Gal4]; MB247-LexA/LexAop2-ACh3.0

Figure 6

UAS-Kir2.1 / CyO; VT064246-Gal4 / TM3

UAS-CsChrimson-mCherry / CyO; VT064246-Gal4 / TM6B

UAS-Trh/UAS-Trh; VT064246-Gal4, Trh01/ VT064246-Gal4, Trh01

UAS-5-HT1a-RNAi/30y-Gal4

QYJ-SI-5HT1a[Gal4]/ QYJ-SI-5HT1a[Gal4]

### DETAILED METHODS

#### Functional imaging

Adult female flies within 2 weeks after eclosion were used for imaging experiments. The fly was mounted to a customized chamber using tape, and a 1 mm X 1 mm rectangular section of tape above the head was removed. The cuticle between the eyes, the air sacs, and the fat bodies were carefully removed in order to expose the brain, which was bathed in adult hemolymph-like solution (AHLS) containing (in mM): 108 NaCl, 5 KCl, 5 HEPES, 5 D-trehalose, 5 sucrose, 26 NaHCO3, 1 NaH2PO4, 2 CaCl2 and 2 MgCl2.

The experiments in Figure 3A-3G were conducted using a Leica SP5 II confocal microscope, with a 488 nm laser for excitation and the 490-560-nm spectrum for the green fluorescence signal. Other functional imaging experiments were conducted using an Olympus FVMPE-RS microscope equipped with a Spectra-Physics InSight X3 two-photon laser, with 920-nm laser for excitation and a 495-540-nm filter to collect the green fluorescence signal. For odorant stimulation, the odorant was diluted 200-fold in mineral oil, then diluted 5-fold in air and delivered to the antenna at a rate of 1000 ml/min. The odorant isoamyl acetate was used for the experiments in Figures 3-4, while 3-octanol (OCT) and 4-methylcyclohexanol (MCH) were used in the experiments in Figures 4-5 and Figure S6-8. For single-photon optogenetic stimulation, a 635-nm laser (Changchun Liangli Photo Electricity Co., Ltd.) was used, and an 18 mW/cm^2^ light was delivered to the brain via an optic fiber. For two-photon optogenetic stimulation, a 1045-nm laser was used, and a 20-mW light was delivered to the region of interest. For electric shock stimulation, two copper wires were attached to the fly’s abdomen and 80-V pulses were delivered. To apply various neurotransmitters (e.g., 5-HT, ACh, DA, OA, Glu, and GABA) and chemicals (e.g., ketanserin, metoclopramide, SB216641, and WAY100635) to the brain, a small patch of the blood-brain-barrier was carefully removed with tweezers before the experiment. The following sampling rates were used: 5 Hz (Figure 3A-3G), 6.8 Hz (Figures 3J-3K, and 4A-4C), 1 Hz (Figures 3L and 4J-4L), 10 Hz (Fig. 4D- 4F), and 4 Hz (Figures 2, 4G-4I and 5).

#### Immunostaining and confocal imaging

The brains of female and male adults within 7-14 days after eclosion were dissected into ice-cold phosphate-buffered saline (PBS), fixed in ice-cold 4% (w/v) paraformaldehyde solution for 1 h, and washed three times with washing buffer (PBS containing 3% NaCl, 1% Triton X-100) for 10 min each. The brains were then incubated in penetration/blocking buffer (PBS containing 2% Triton X- 100 and 10% normal goat serum) for 20 h at 4°C on a shaker. The brains were then incubated with primary antibodies (diluted in PBS containing 0.25% Triton X-100 and 1% normal goat serum) for 24 hours at 4°C, and then washed three times in washing buffer for 10 min each on a shaker. The brains were then incubated with the appropriate secondary antibodies (diluted in PBS containing 0.25% Triton X-100 and 1% normal goat serum) overnight at 4°C in the dark, then washed three times with washing buffer for 10 min each on a shaker. The samples were mounted with Fluoroshield and kept in the dark. The following antibodies were used at the indicated dilutions: chicken anti-GFP (1:500), rabbit anti-mCherry (1:500), mouse anti-nc82 (1:40), Alexa Fluor 488 goat anti-chicken (1:500), Alex Fluor 555 goat anti-rabbit (1:500), and Alex Fluor 647 goat anti-mouse (1:500). Fluorescence images were obtained using a Nikon Ti-E A1 confocal microscope. Alexa Fluor 488, Alexa Fluor 555, and Alexa Fluor 647 were excited using a 485-nm, 559-nm, and 638-nm laser, respectively, and imaged using a 525/50-nm, 595/50-nm, and 700/75-nm filter, respectively.

#### Behavioral assay

These experiments were performed in a dark room at 22°C with 50-60% humidity. Flies within 24- 72 hours after eclosion were transferred to a new tube 12 hours before the experiment. The airflow rates of the training arm and the testing arms were maintained at 800 ml/min throughout the experiment. Before training, 50-100 flies were loaded in the training arm and accommodated for 2 min. During training, the CS+ (diluted by 67-fold in mineral oil) was delivered via the airflow for 10 s. Three 90-V electric shocks were delivered via the copper grid contained within the training arm at 0.2 Hz, with a varying ISI. For optogenetic stimulation, a 635-nm laser (Changchun Liangli Photo Electricity Co., Ltd.) was used, and a 10 mW/cm^2^ light was delivered to the training arm via an optic fiber. 2 min after the end of CS+, the CS- (diluted by 67-fold in mineral oil) was delivered via the airflow for 10 s. One min after training, the flies were transferred to the elevator and allowed to accommodate for 3 min before testing. During testing, the paired and unpaired conditioned stimuli (CS+ and CS-, respectively) were delivered from two ends of the arms for 30 s, after which the number of flies in each arm (N) was counted. The performance index was calculated using the following formula: [N (CS+) – N (CS-)] / [N (CS+) + N (CS-)]. One group of flies were used in only one trial training and testing. To reduce the possible bias of innate preference, each data point is the average result of two groups of flies (electric shock paired with OCT in one group, and electric shock paired with MCH in the other group).

### Quantification and data analysis

Imaging data from *Drosophila* brains were firstly processed using Image J software (National Institutes of Health), followed by replotting graphs using Origin 9.1 (OriginLab). The fluorescence responses (ΔF/F0) were calculated using the formula (F-F0)/F0, in which F0 is the basal fluorescent signal. The Relative ∫ ΔF/F0 (Figure 2 and 5) was the calculation of the area under curve during odor application followed by normalization to that in control group. The behavioral performance index (Figure 1 and 6) was calculated as mentioned above in behavioral assay part. For better comparison, in the sigmoid function fitted traces of learning behavior, the performance index against ISI = 5 s was related to 1. In the sigmoid function fitted traces for synaptic plasticity, the ΔACh is the ∫ ΔF/F0 (Pre) - ∫ ΔF/F0 (Post).

Except where indicated otherwise, all summary data were presented as the Mean ± SEM, and group differences were analyzed using Student’s t-test and One-Way ANOVA test.

## Acknowledgments

We thank Y.R. (Peking University) and State Key Laboratory of Membrane Biology for providing support regarding the two-photon microscope, and Y.L. (Institute of Biophysics, Chinese Academy of Science) for sharing the confocal microscope. We thank L.Luo, J.Wang, Q.Gaudry, S.Owen, S.Tomchik, A.Lutas, R.Yasuda, L.Liang, R.Davis, L.Liu, Y.Zhong, J.Ren, P.Fan, S.Zhang, B.Zhao, B.Deng, F.Wang and K.Wang for valuable feedback of the manuscript.

## Author Contributions

Y.L. conceived and supervised the project. J.Z. and X.L. performed the immunofluorescence imaging and all functional imaging experiments, unless otherwise noted. Z.Z., X.L., and J.Z. performed the behavioral experiments and analyzed the EM data. X.L. analyzed the transcriptomics data. M.L. performed the neurotransmitter perfusion experiments. Y.W. contributed to the experiments using hM4Di. K.T. and Y.W. contributed to the synaptic plasticity experiments.

X.X. contributed to the fly preparation. J.W. and M.J. provided the 5-HT1.0 and ACh3.0 sensors, respectively. All authors contributed to the data interpretation and data analysis. Y.L. wrote the manuscript with input from all other authors.

---

**Figure S1.**
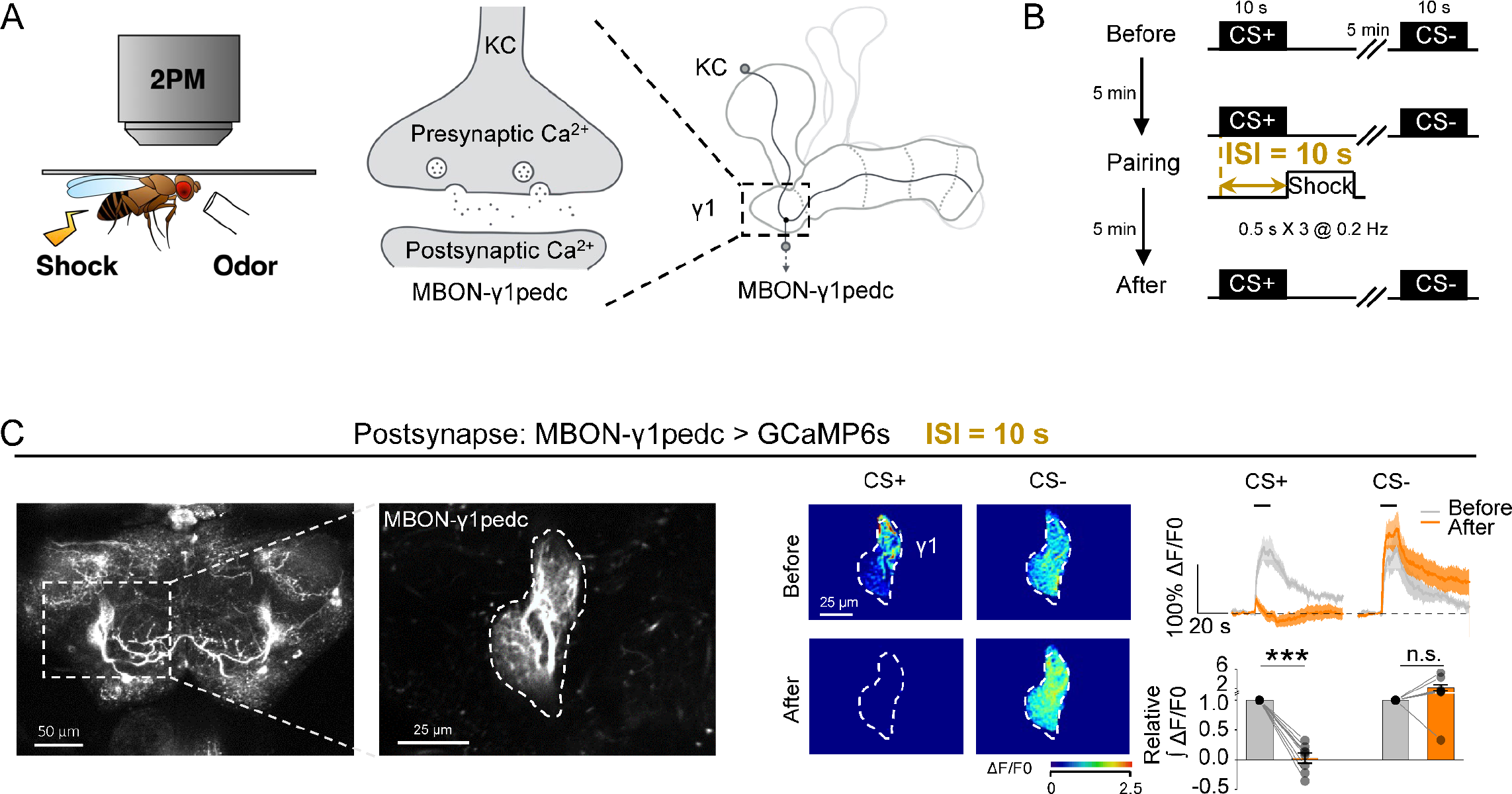
Ca^2+^ signals reveal changes in synaptic plasticity in the γ1 compartment. (A) Schematic diagram depicting the strategy used to image Ca**2+** induced by odorant application or electric shock. signals in the MBON-γ1pedc (B) The experimental protocol. CS+ and CS- represent the paired conditioned stimulus and the unpaired conditioned stimulus, respectively. (C) Fluorescence images (left), change in GCaMP6s fluorescence (middle), average traces (top right), and relative group responses (bottom right) of postsynaptic Ca**2+** signals in response to CS+ and CS- before and after pairing. \*\*\**p*<0.001 and n.s., not significant (Student’s *t*-test).

**Figure S2.**
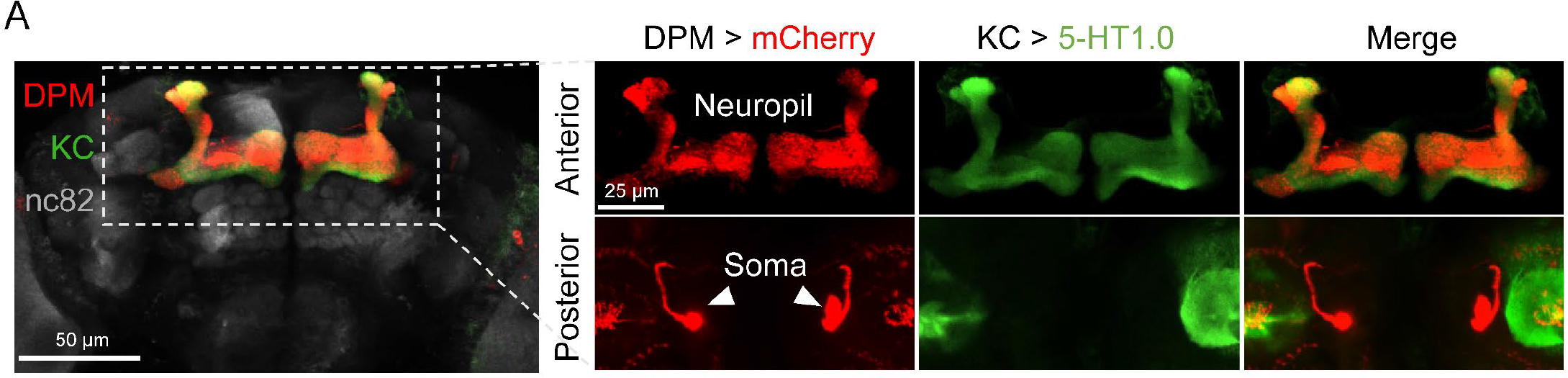
Immunofluorescence images of the DPM neuron and KCs. Immunofluorescence images of the dissected brain from a fly expressing mCherry (red) in the DPM neuron and 5-HT1.0 (green) in the KCs. Each image is a projection of several slices through the MB. Arrowheads indicate the somas of the two DPM neurons.

**Figure S3.**
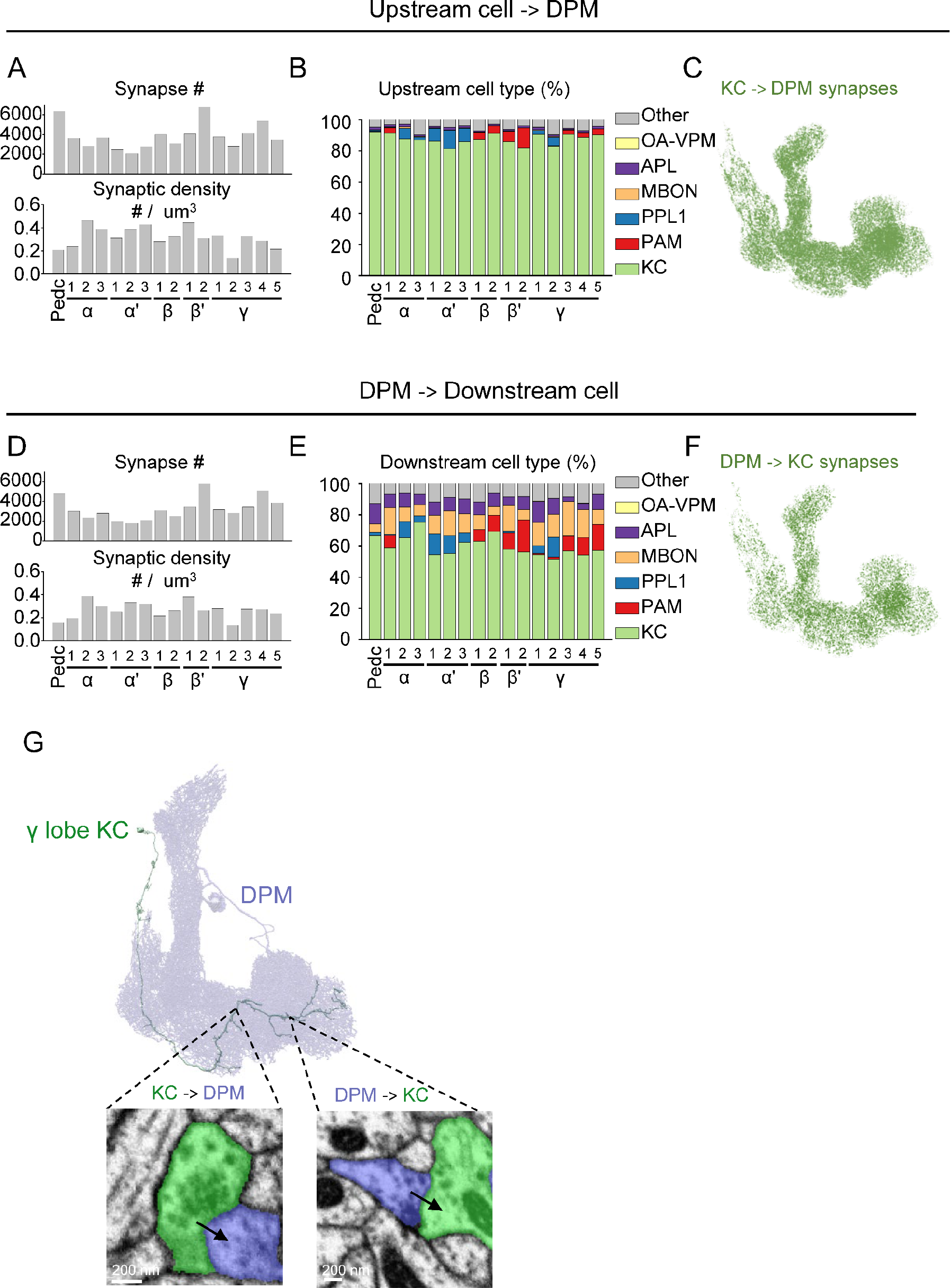
EM connectomics reveals reciprocal connections between the DPM neuron and KCs. (**A** and **B**) Quantification of the number (**A**, top) and density (**A**, bottom) of synapses upstream from the DPM, and percentage of cell types in the indicated MB compartments. (C) Synapses from the KCs to the DPM neuron. (**D-F**) Similar to (**A**-**C**), except that the synapses downstream of the DPM were measured. (**G**) Representative cartoon and EM images of a KC forming reciprocal connections with the DPM neuron in the γ lobe. Arrows indicate the orientation of the annotated synapses. Version 1.1 of the hemibrain connectome (Scheffer et al., 2020) was used for the analysis, and only synapses with a confidence value >0.75 were included. Pedc, peduncle; OA-VPM, octopaminergic VPM neurons; APL, GABAergic anterior paired lateral neurons; MBON, mushroom body output neurons; PPL1, paired posterior lateral 1 cluster neurons; PAM, protocerebral anterior medial cluster neurons; KC, Kenyon cell.

**Figure S4.**
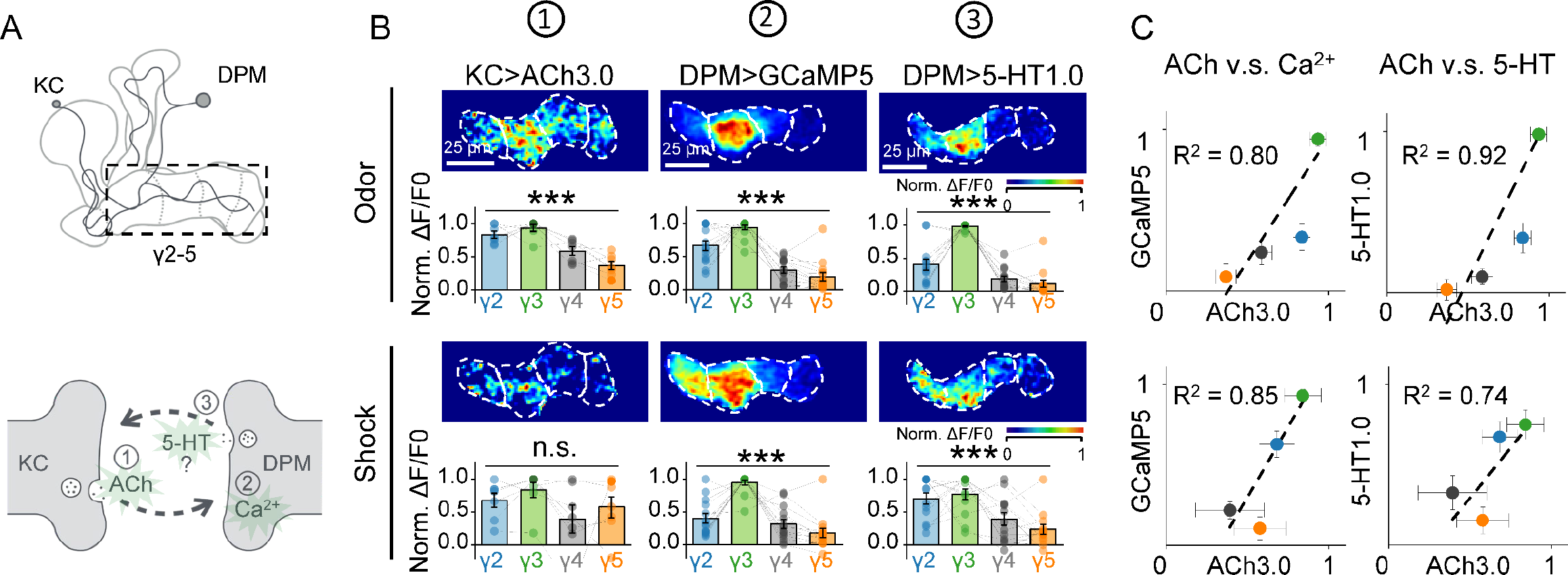
The heterogenous pattern of 5-HT release is highly correlated with the ACh release from KCs. (A) Schematic diagram depicting the strategy used to image ACh, 5-HT, and Ca^2+^in the γ2-5 compartments. (B) Representative normalized pseudocolor images and group data of the indicated fluorescence signals measured in the γ2-5 compartments in response to a 1-s odorant stimulation or a 0.5-s electric shock. For each fly, fluorescence signals were normalized to the compartment with the highest response. (C) Correlation analysis of the change in fluorescence measured in response to the indicated stimuli. The data were fit to a linear function, with the corresponding correlation coefficients shown. Group data are presented as the mean ± SEM, overlaid with the data obtained from each fly. \**p*<0.05, (One-way ANOVA).

**Figure S5.**
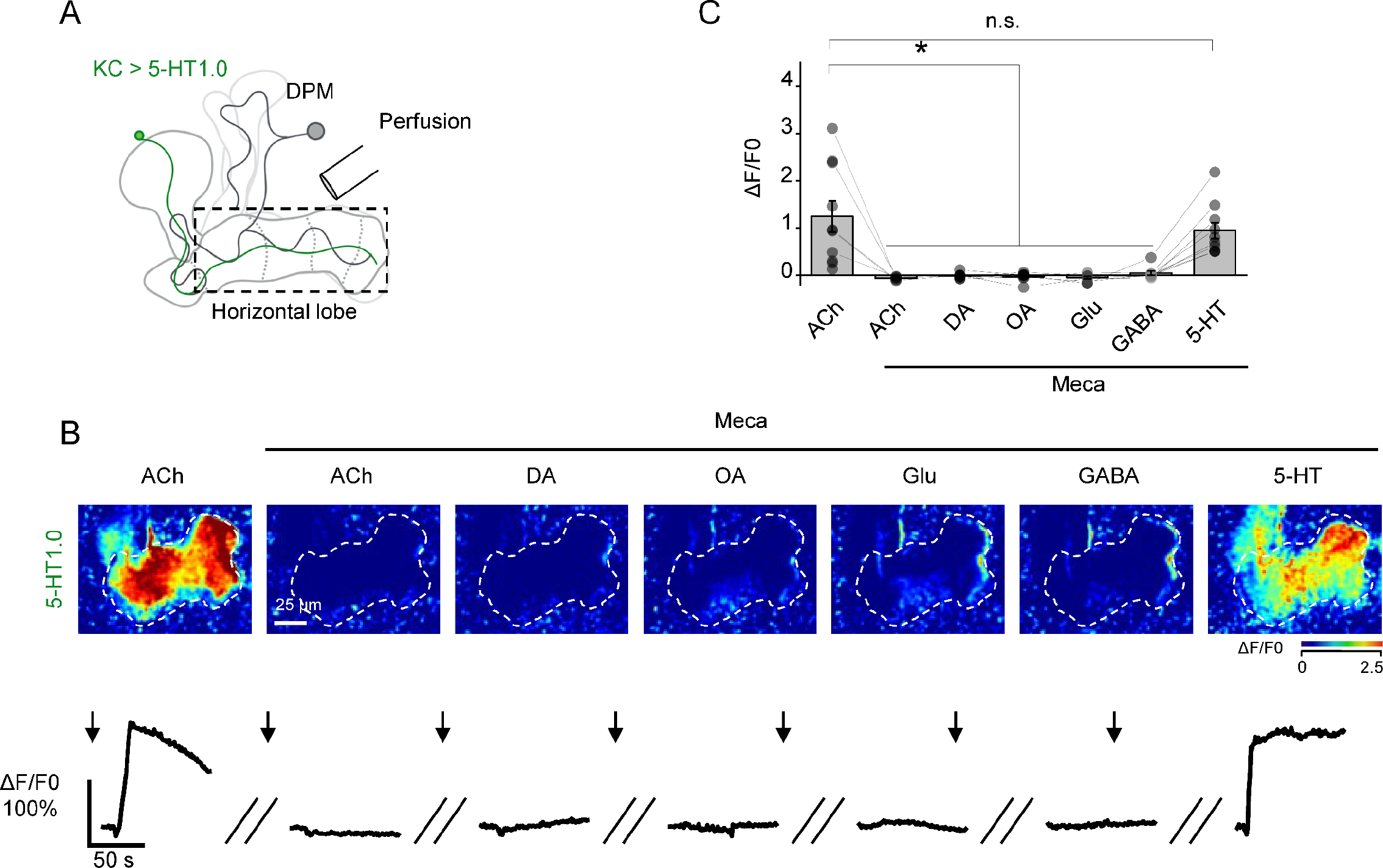
ACh application induces 5-HT release via nAChRs. (**A**) Schematic diagram depicting the strategy used for the perfusion experiments; 5-HT was measured using 5-HT1.0 expressed in KCs. (**B** and **C**) Representative pseudocolor images (**B**, top), corresponding traces (**B**, bottom), and group data (**C**) of the change in 5-HT1.0 fluorescence in response to application of the indicated neurotransmitters (at 1 mM) in the absence or presence of the nicotinic ACh receptor antagonist Meca (100 μM). \**p*<0.05 and n.s., not significant (Student’s *t*-test). ACh, acetylcholine; DA, dopamine; OA, octopamine; Glu, glutamate; GABA, gamma-aminobutyric acid.

**Figure S6.**
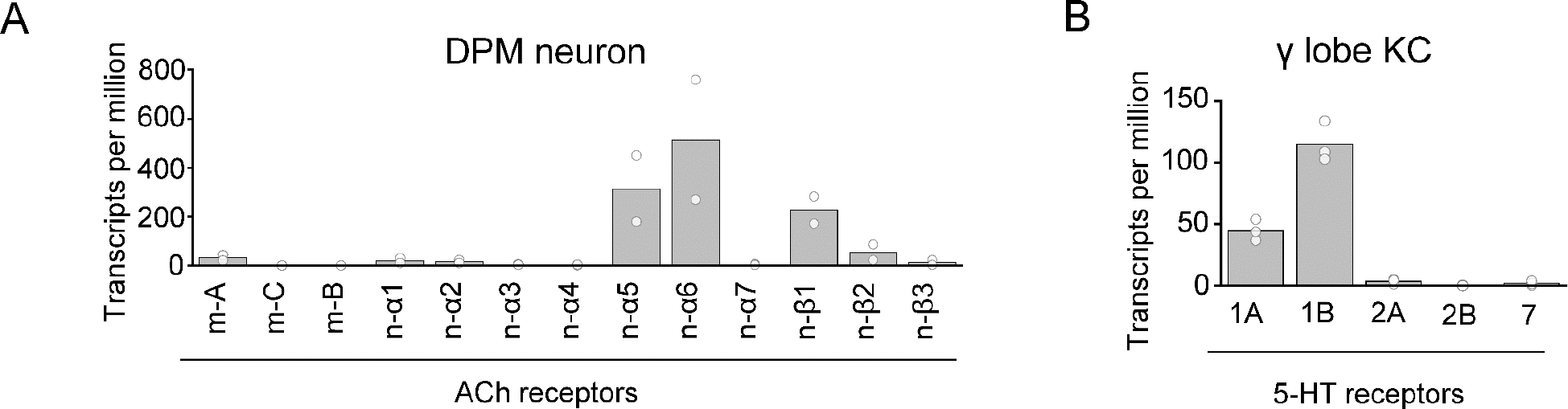
Transcriptomics analysis of ACh receptor subtypes and 5-HT receptor subtypes in the DPM neuron and KCs, respectively. (A) Relative abundance of the indicated transcripts measured in DPM neurons. (B) Relative abundance of the indicated transcripts measured in KCs in the γ lobe. Group data are shown as the mean value overlaid with data from each sample. One sample includes 123 or 130 cells (**a**), or 2500 cells (**B**), collected from 60-100 fly brains. The transcript database (Aso et al., 2019) was used for analysis.

**Figure S7.**
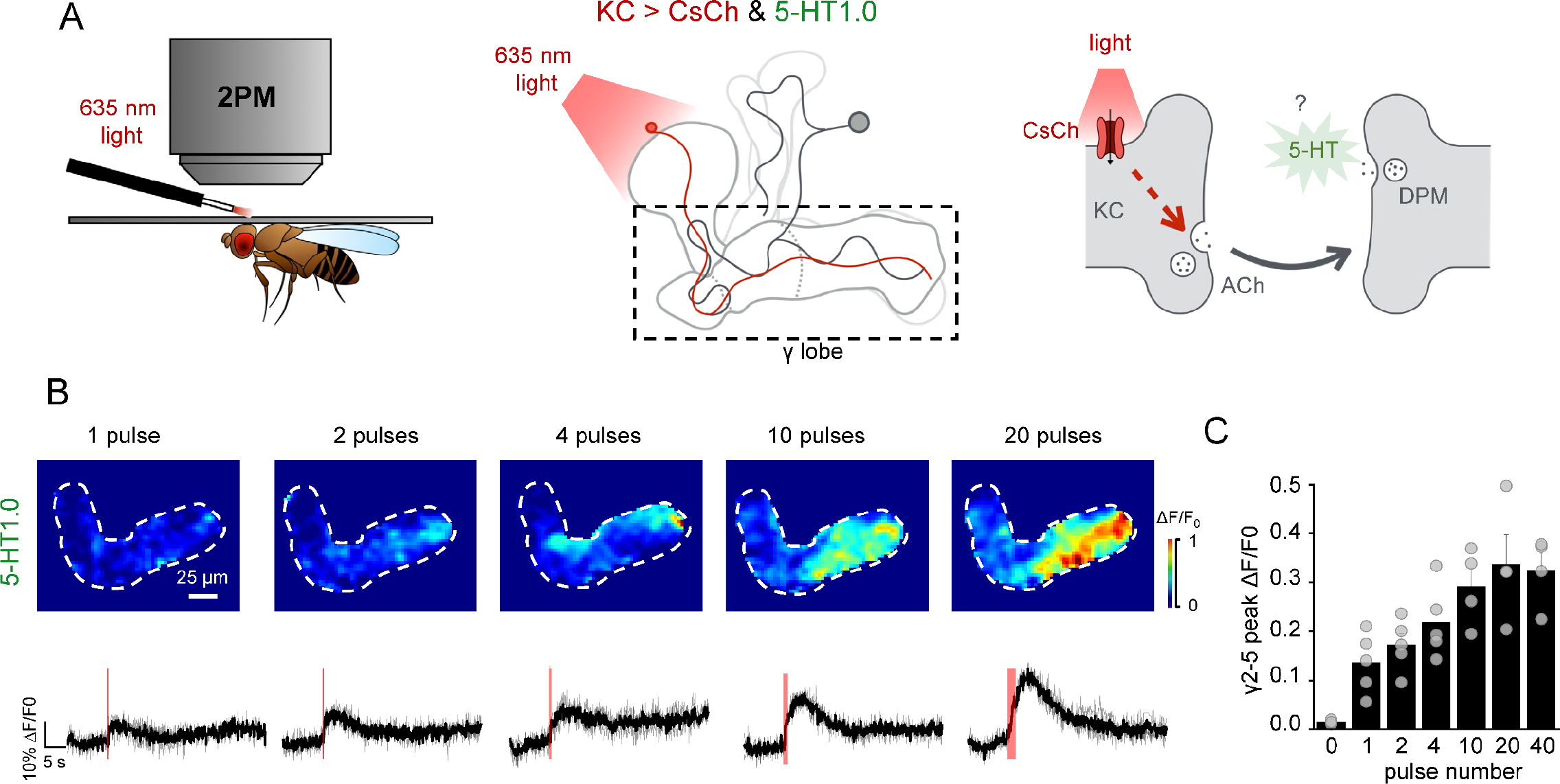
DPM receive excitatory input from KCs. (**A**) Schematic diagram depicting the strategy used for the experiment. KCs were activated by 635 nm light (10Hz, 1ms/pulse) with CsChrimson. 5-HT was measured using 5-HT1.0 expressed in KCs. (**B** and **C**) Representative pseudocolor images (**B**) and group data (**C**) of the change in 5-HT1.0 fluorescence in response to different pulses activation of KCs.

**Figure S8.**
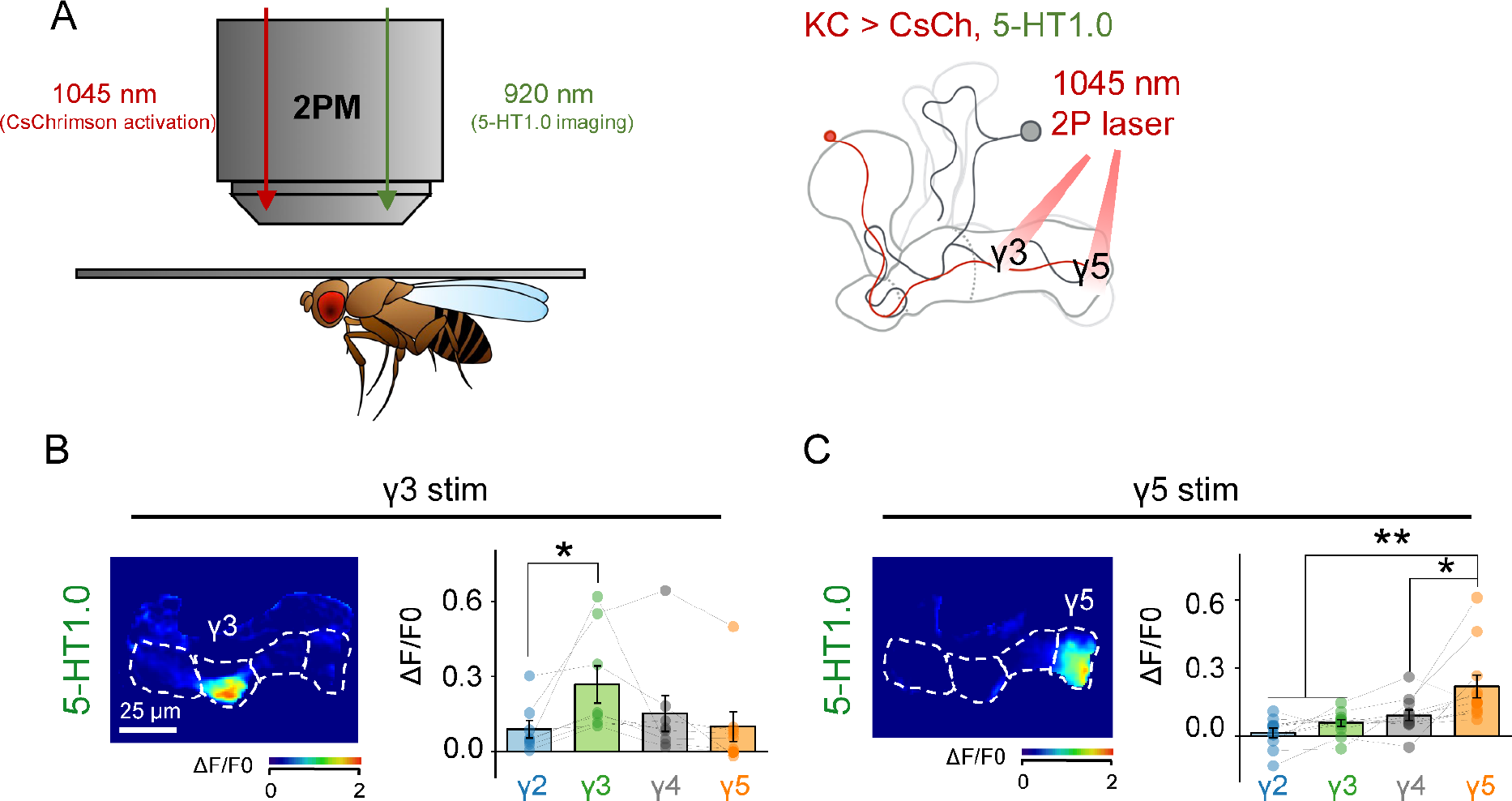
Local activation of KCs induces heterogenous 5-HT release. (**A**) Schematic diagram depicting the strategy used for the experiment. A 1045-nm two-photon laser was used to locally activate CsChrimson expressed in KCs. 5-HT signal was measured with 5-HT1.0 expressed in KCs. (**B** and **C**) Representative pseudocolor images (left) and group data (right) of the change in 5- HT1.0 fluorescence in response to local optogenetic stimulation in the γ3 (**B**) and γ5 (**C**) compartments. \**p*<0.05, \*\**p*<0.01, and n.s., not significant (Student’s *t*-test)

**Figure S9.**
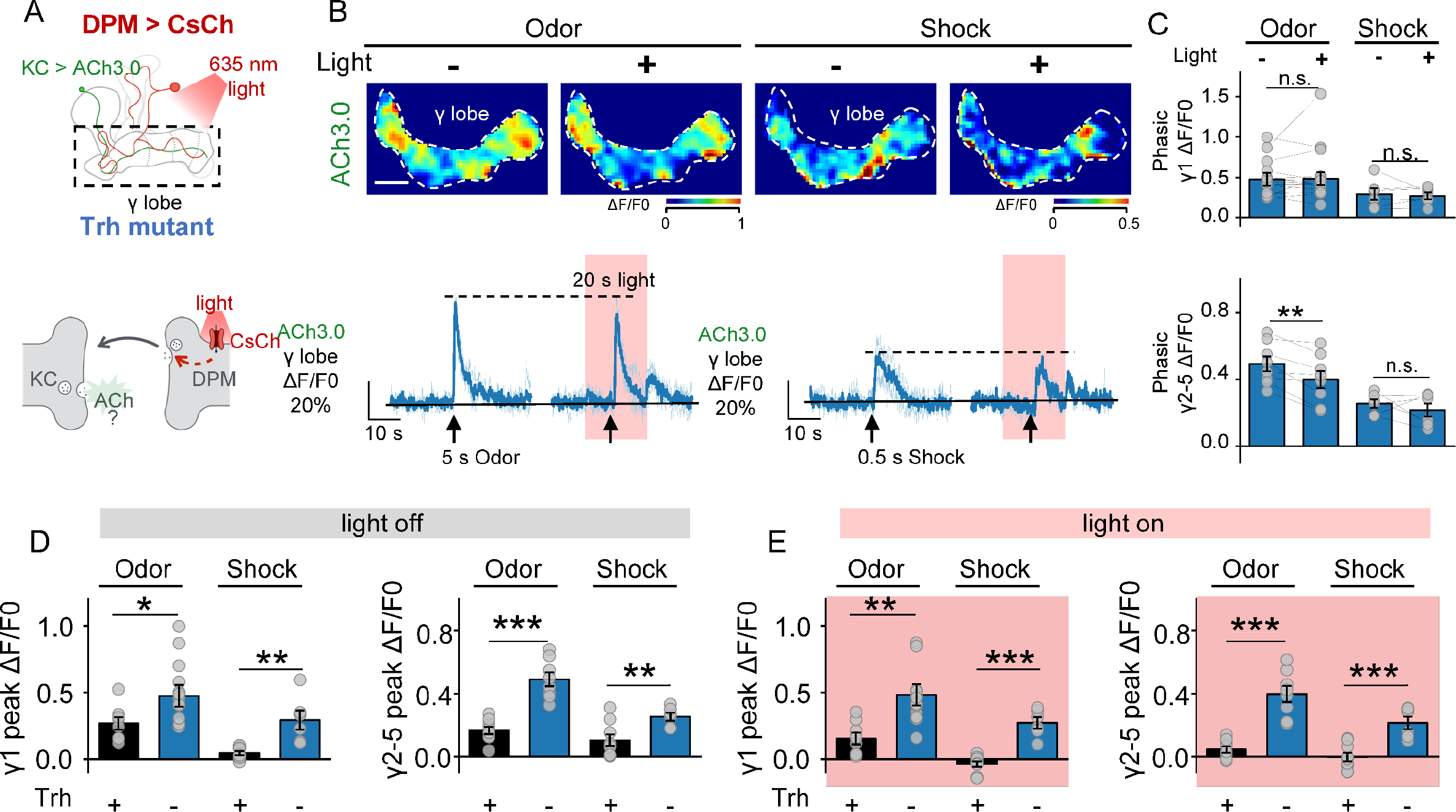
5-HT from DPM provides feedback inhibition to KCs. (**A**) Schematic diagram depicting the experimental setup for the subsequent experiments. In Trh mutant flies, DPM is activated with CsChrimson by 635-nm light at 10Hz, 1 ms / pulse. ACh signals are measured with ACh3.0 expressed in KCs. (**B** and **C**) Representative pseudocolor images B, top), traces (**B**, bottom), and group data (**C**) of the change in ACh3.0 fluorescence in response to a 20-s optogenetic stimulation in saline. (**D** and **E**) Group comparison of odor and shock evoked ACh release in control flies (black) and Trh mutant flies (blue) without (**D**) or with DPM activation (**E**). Data plotted with Meas ± SEM. \**p*<0.05, \*\**p*<0.01, and \*\*\**p*<0.001 (Student’s *t*-test).

## References

Akalal, D.B., Yu, D., and Davis, R.L. (2010). A late-phase, long-term memory trace forms in the gamma neurons of Drosophila mushroom bodies after olfactory classical conditioning. J Neurosci 30, 16699–16708.

Amin, H., Apostolopoulou, A.A., Suarez-Grimalt, R., Vrontou, E., and Lin, A.C. (2020). Localized inhibition in the Drosophila mushroom body. Elife 9.

Armbruster, B.N., Li, X., Pausch, M.H., Herlitze, S., and Roth, B.L. (2007). Evolving the lock to fit the key to create a family of G protein-coupled receptors potently activated by an inert ligand. Proc Natl Acad Sci U S A 104, 5163–5168.

Aso, Y., Hattori, D., Yu, Y., Johnston, R.M., Iyer, N.A., Ngo, T.T., Dionne, H., Abbott, L.F., Axel, R., Tanimoto, H., et al. (2014). The neuronal architecture of the mushroom body provides a logic for associative learning. Elife 3, e04577.

Aso, Y., Ray, R.P., Long, X., Bushey, D., Cichewicz, K., Ngo, T.T., Sharp, B., Christoforou, C., Hu, A., Lemire, A.L., et al. (2019). Nitric oxide acts as a cotransmitter in a subset of dopaminergic neurons to diversify memory dynamics. Elife 8.

Aso, Y., and Rubin, G.M. (2016). Dopaminergic neurons write and update memories with cell-type-specific rules. Elife 5.

Barnstedt, O., Owald, D., Felsenberg, J., Brain, R., Moszynski, J.P., Talbot, C.B., Perrat, P.N., and Waddell, S. (2016). Memory-Relevant Mushroom Body Output Synapses Are Cholinergic. Neuron 89, 1237–1247.

Becnel, J., Johnson, O., Majeed, Z.R., Tran, V., Yu, B., Roth, B.L., Cooper, R.L., Kerut, E.K., and Nichols, C.D. (2013). DREADDs in Drosophila: a pharmacogenetic approach for controlling behavior, neuronal signaling, and physiology in the fly. Cell Rep 4, 1049–1059.

Bernstein, A.L. (1934). Temporal Factors in the Formation of Conditioned Eyelid Reactions in Human Subjects. The Journal of General Psychology.

Berry, J.A., Cervantes-Sandoval, I., Nicholas, E.P., and Davis, R.L. (2012). Dopamine is required for learning and forgetting in Drosophila. Neuron 74, 530–542.

Berry, J.A., Phan, A., and Davis, R.L. (2018). Dopamine Neurons Mediate Learning and Forgetting through Bidirectional Modulation of a Memory Trace. Cell Rep 25, 651–662 e655.

Bilz, F., Geurten, B.R.H., Hancock, C.E., Widmann, A., and Fiala, A. (2020). Visualization of a Distributed Synaptic Memory Code in the Drosophila Brain. Neuron 106, 963–976 e964.

Bolbecker, A.R., Steinmetz, A.B., Mehta, C.S., Forsyth, J.K., Klaunig, M.J., Lazar, E.K., Steinmetz, J.E., O’Donnell, B.F., and Hetrick, W.P. (2011). Exploration of cerebellar- dependent associative learning in schizophrenia: effects of varying and shifting interstimulus interval on eyeblink conditioning. Behav Neurosci 125, 687–698.

Boto, T., Louis, T., Jindachomthong, K., Jalink, K., and Tomchik, S.M. (2014). Dopaminergic modulation of cAMP drives nonlinear plasticity across the Drosophila mushroom body lobes. Curr Biol 24, 822–831.

Boto, T., Stahl, A., Zhang, X., Louis, T., and Tomchik, S.M. (2019). Independent Contributions of Discrete Dopaminergic Circuits to Cellular Plasticity, Memory Strength, and Valence in Drosophila. Cell Rep 27, 2014–2021 e2012.

Bouzaiane, E., Trannoy, S., Scheunemann, L., Placais, P.Y., and Preat, T. (2015). Two independent mushroom body output circuits retrieve the six discrete components of Drosophila aversive memory. Cell Rep 11, 1280–1292.

Brzosko, Z., Mierau, S.B., and Paulsen, O. (2019). Neuromodulation of Spike-Timing- Dependent Plasticity: Past, Present, and Future. Neuron 103, 563–581.

Buhot, M.C., Martin, S., and Segu, L. (2000). Role of serotonin in memory impairment. Ann Med 32, 210–221.

Burke, C.J., Huetteroth, W., Owald, D., Perisse, E., Krashes, M.J., Das, G., Gohl, D., Silies, M., Certel, S., and Waddell, S. (2012). Layered reward signalling through octopamine and dopamine in Drosophila. Nature 492, 433–437.

Carew, T.J., Walters, E.T., and Kandel, E.R. (1981). Classical conditioning in a simple withdrawal reflex in Aplysia californica. J Neurosci 1, 1426–1437.

Claridge-Chang, A., Roorda, R.D., Vrontou, E., Sjulson, L., Li, H., Hirsh, J., and Miesenbock, G. (2009). Writing memories with light-addressable reinforcement circuitry. Cell 139, 405–415.

Coates, K.E., Calle-Schuler, S.A., Helmick, L.M., Knotts, V.L., Martik, B.N., Salman, F., Warner, L.T., Valla, S.V., Bock, D.D., and Dacks, A.M. (2020). The Wiring Logic of an Identified Serotonergic Neuron That Spans Sensory Networks. J Neurosci 40, 6309–6327.

Coates, K.E., Majot, A.T., Zhang, X., Michael, C.T., Spitzer, S.L., Gaudry, Q., and Dacks, A.M. (2017). Identified Serotonergic Modulatory Neurons Have Heterogeneous Synaptic Connectivity within the Olfactory System of Drosophila. J Neurosci 37, 7318–7331.

Cohn, R., Morantte, I., and Ruta, V. (2015). Coordinated and Compartmentalized Neuromodulation Shapes Sensory Processing in Drosophila. Cell 163, 1742–1755.

Connors, B.W. (2012). Tales of a dirty drug: carbenoxolone, gap junctions, and seizures. Epilepsy Curr 12, 66–68.

Croset, V., Treiber, C.D., and Waddell, S. (2018). Cellular diversity in the Drosophila midbrain revealed by single-cell transcriptomics. Elife 7.

Dacks, A.M., Christensen, T.A., and Hildebrand, J.G. (2006). Phylogeny of a serotonin- immunoreactive neuron in the primary olfactory center of the insect brain. J Comp Neurol 498, 727–746.

Davis, R.L., Cherry, J., Dauwalder, B., Han, P.L., and Skoulakis, E. (1995). The cyclic AMP system and Drosophila learning. Mol Cell Biochem 149-150, 271-278.

Dudai, Y., Jan, Y.N., Byers, D., Quinn, W.G., and Benzer, S. (1976). dunce, a mutant of Drosophila deficient in learning. Proc Natl Acad Sci U S A 73, 1684–1688.

Dudai, Y., Sher, B., Segal, D., and Yovell, Y. (1985). Defective responsiveness of adenylate cyclase to forskolin in the Drosophila memory mutant rutabaga. J Neurogenet 2, 365–380.

Dylla, K.V., Raiser, G., Galizia, C.G., and Szyszka, P. (2017). Trace Conditioning in Drosophila Induces Associative Plasticity in Mushroom Body Kenyon Cells and Dopaminergic Neurons. Front Neural Circuits 11, 42.

Eichler, K., Li, F., Litwin-Kumar, A., Park, Y., Andrade, I., Schneider-Mizell, C.M., Saumweber, T., Huser, A., Eschbach, C., Gerber, B., et al. (2017). The complete connectome of a learning and memory centre in an insect brain. Nature 548, 175–182.

Felsenberg, J., Barnstedt, O., Cognigni, P., Lin, S., and Waddell, S. (2017). Re-evaluation of learned information in Drosophila. Nature 544, 240–244.

Felsenberg, J., Jacob, P.F., Walker, T., Barnstedt, O., Edmondson-Stait, A.J., Pleijzier, M.W., Otto, N., Schlegel, P., Sharifi, N., Perisse, E., et al. (2018). Integration of Parallel Opposing Memories Underlies Memory Extinction. Cell 175, 709–722 e715.

Fonseca, M.S., Murakami, M., and Mainen, Z.F. (2015). Activation of dorsal raphe serotonergic neurons promotes waiting but is not reinforcing. Curr Biol 25, 306–315.

Frings, M., Gaertner, K., Buderath, P., Gerwig, M., Christiansen, H., Schoch, B., Gizewski, E.R., Hebebrand, J., and Timmann, D. (2010). Timing of conditioned eyeblink responses is impaired in children with attention-deficit/hyperactivity disorder. Exp Brain Res 201, 167–176.

Ganguly, A., Qi, C., Bajaj, J., and Lee, D. (2020). Serotonin receptor 5-HT7 in Drosophila mushroom body neurons mediates larval appetitive olfactory learning. Sci Rep 10, 21267.

Gerber, B., König, C., Fendt, M., Andreatta, M., Romanos, M., Pauli, P., and Yarali, A. (2019). Timing-dependent valence reversal: a principle of reinforcement processing and its possible implications. Current Opinion in Behavioral Sciences 26, 114–120.

Gerber, B., Yarali, A., Diegelmann, S., Wotjak, C.T., Pauli, P., and Fendt, M. (2014). Pain- relief learning in flies, rats, and man: basic research and applied perspectives. Learn Mem 21, 232–252.

Gervasi, N., Tchenio, P., and Preat, T. (2010). PKA dynamics in a Drosophila learning center: coincidence detection by rutabaga adenylyl cyclase and spatial regulation by dunce phosphodiesterase. Neuron 65, 516–529.

Handler, A., Graham, T.G.W., Cohn, R., Morantte, I., Siliciano, A.F., Zeng, J., Li, Y., and Ruta, V. (2019). Distinct Dopamine Receptor Pathways Underlie the Temporal Sensitivity of Associative Learning. Cell 178, 60–75 e19.

Harmer, C.J., Bhagwagar, Z., Cowen, P.J., and Goodwin, G.M. (2002). Acute administration of citalopram facilitates memory consolidation in healthy volunteers. Psychopharmacology (Berl) 163, 106–110.

Harvey, J.A. (2003). Role of the serotonin 5-HT2A receptor in learning. Learn Memory 10, 355–362.

Harvey, J.A., Gormezano, I., Cool-Hauser, V.A., and Schindler, C.W. (1988). Effects of LSD on classical conditioning as a function of CS-UCS interval: relationship to reflex facilitation. Pharmacol Biochem Behav 30, 433–441.

Hawkins, R.D., Carew, T.J., and Kandel, E.R. (1986). Effects of interstimulus interval and contingency on classical conditioning of the Aplysia siphon withdrawal reflex. J Neurosci 6, 1695–1701.

Haynes, P.R., Christmann, B.L., and Griffith, L.C. (2015). A single pair of neurons links sleep to memory consolidation in Drosophila melanogaster. Elife 4.

Heisenberg, M. (2003). Mushroom body memoir: from maps to models. Nat Rev Neurosci 4, 266–275.

Hige, T. (2018). What can tiny mushrooms in fruit flies tell us about learning and memory? Neurosci Res 129, 8–16.

Hige, T., Aso, Y., Modi, M.N., Rubin, G.M., and Turner, G.C. (2015). Heterosynaptic Plasticity Underlies Aversive Olfactory Learning in Drosophila. Neuron 88, 985–998.

Himmelreich, S., Masuho, I., Berry, J.A., MacMullen, C., Skamangas, N.K., Martemyanov, K.A., and Davis, R.L. (2017). Dopamine Receptor DAMB Signals via Gq to Mediate Forgetting in Drosophila. Cell Rep 21, 2074–2081.

Inada, K., Tsuchimoto, Y., and Kazama, H. (2017). Origins of Cell-Type-Specific Olfactory Processing in the Drosophila Mushroom Body Circuit. Neuron 95, 357–367 e354.

Jing, M., Li, Y., Zeng, J., Huang, P., Skirzewski, M., Kljakic, O., Peng, W., Qian, T., Tan, K., Zou, J., et al. (2020). An optimized acetylcholine sensor for monitoring in vivo cholinergic activity. Nat Methods 17, 1139–1146.

Jing, M., Zhang, P., Wang, G., Feng, J., Mesik, L., Zeng, J., Jiang, H., Wang, S., Looby, J.C., Guagliardo, N.A., et al. (2018). A genetically encoded fluorescent acetylcholine indicator for in vitro and in vivo studies. Nat Biotechnol 36, 726–737.

Johnson, O., Becnel, J., and Nichols, C.D. (2011). Serotonin receptor activity is necessary for olfactory learning and memory in Drosophila melanogaster. Neuroscience 192, 372–381.

Kandel, E.R. (2001). The molecular biology of memory storage: a dialogue between genes and synapses. Science 294, 1030–1038.

Kandel, E.R., and Schwartz, J.H. (1982). Molecular biology of learning: modulation of transmitter release. Science 218, 433–443.

Keene, A.C., Krashes, M.J., Leung, B., Bernard, J.A., and Waddell, S. (2006). Drosophila dorsal paired medial neurons provide a general mechanism for memory consolidation. Curr Biol 16, 1524–1530.

Keene, A.C., Stratmann, M., Keller, A., Perrat, P.N., Vosshall, L.B., and Waddell, S. (2004). Diverse odor-conditioned memories require uniquely timed dorsal paired medial neuron output. Neuron 44, 521–533.

Kim, Y.C., Lee, H.G., and Han, K.A. (2007). D1 dopamine receptor dDA1 is required in the mushroom body neurons for aversive and appetitive learning in Drosophila. J Neurosci 27, 7640–7647.

Krashes, M.J., Keene, A.C., Leung, B., Armstrong, J.D., and Waddell, S. (2007). Sequential use of mushroom body neuron subsets during drosophila odor memory processing. Neuron 53, 103–115.

Lee, P.T., Lin, H.W., Chang, Y.H., Fu, T.F., Dubnau, J., Hirsh, J., Lee, T., and Chiang, A.S. (2011). Serotonin-mushroom body circuit modulating the formation of anesthesia-resistant memory in Drosophila. Proc Natl Acad Sci U S A 108, 13794–13799.

Lei, Z., Chen, K., Li, H., Liu, H., and Guo, A. (2013). The GABA system regulates the sparse coding of odors in the mushroom bodies of Drosophila. Biochem Biophys Res Commun 436, 35–40.

Levin, L.R., Han, P.L., Hwang, P.M., Feinstein, P.G., Davis, R.L., and Reed, R.R. (1992). The Drosophila learning and memory gene rutabaga encodes a Ca2+/Calmodulin-responsive adenylyl cyclase. Cell 68, 479–489.

Li, F., Lindsey, J.W., Marin, E.C., Otto, N., Dreher, M., Dempsey, G., Stark, I., Bates, A.S., Pleijzier, M.W., Schlegel, P., et al. (2020). The connectome of the adult Drosophila mushroom body provides insights into function. Elife 9.

Li, Y., Zhong, W., Wang, D., Feng, Q., Liu, Z., Zhou, J., Jia, C., Hu, F., Zeng, J., Guo, Q., et al. (2016). Serotonin neurons in the dorsal raphe nucleus encode reward signals. Nat Commun 7, 10503.

Lin, A.C., Bygrave, A.M., de Calignon, A., Lee, T., and Miesenbock, G. (2014). Sparse, decorrelated odor coding in the mushroom body enhances learned odor discrimination. Nat Neurosci 17, 559–568.

Liu, C., Placais, P.Y., Yamagata, N., Pfeiffer, B.D., Aso, Y., Friedrich, A.B., Siwanowicz, I., Rubin, G.M., Preat, T., and Tanimoto, H. (2012). A subset of dopamine neurons signals reward for odour memory in Drosophila. Nature 488, 512–516.

Liu, X., and Davis, R.L. (2009). The GABAergic anterior paired lateral neuron suppresses and is suppressed by olfactory learning. Nat Neurosci 12, 53–59.

Liu, Y.H., Smith, S.J., Mihalas, S., Shea-Brown, E., and Sumbul, U. (2020a). A solution to temporal credit assignment using cell-type-specific modulatory signals. bioRxiv.

Liu, Z., Lin, R., and Luo, M. (2020b). Reward Contributions to Serotonergic Functions. Annu Rev Neurosci 43, 141–162.

Liu, Z., Zhou, J., Li, Y., Hu, F., Lu, Y., Ma, M., Feng, Q., Zhang, J.E., Wang, D., Zeng, J., et al. (2014). Dorsal raphe neurons signal reward through 5-HT and glutamate. Neuron 81, 1360–1374.

Livingstone, M.S., Sziber, P.P., and Quinn, W.G. (1984). Loss of calcium/calmodulin responsiveness in adenylate cyclase of rutabaga, a Drosophila learning mutant. Cell 37, 205–215.

Lottem, E., Banerjee, D., Vertechi, P., Sarra, D., Lohuis, M.O., and Mainen, Z.F. (2018). Activation of serotonin neurons promotes active persistence in a probabilistic foraging task. Nat Commun 9, 1000.

Louis, T., Stahl, A., Boto, T., and Tomchik, S.M. (2018). Cyclic AMP-dependent plasticity underlies rapid changes in odor coding associated with reward learning. Proc Natl Acad Sci U S A 115, E448–E457.

Mao, Z., and Davis, R.L. (2009). Eight different types of dopaminergic neurons innervate the Drosophila mushroom body neuropil: anatomical and physiological heterogeneity. Front Neural Circuits 3, 5.

McAllister, W.R. (1953). Eyelid conditioning as a function of the CS-US interval. J Exp Psychol 45, 417–422.

McCurdy, L.Y., Sareen, P., Davoudian, P.A., and Nitabach, M.N. (2021). Dopaminergic mechanism underlying reward-encoding of punishment omission during reversal learning in Drosophila. Nat Commun 12, 1115.

McGlinchey-Berroth, R., Brawn, C., and Disterhoft, J.F. (1999). Temporal discrimination learning in severe amnesic patients reveals an alteration in the timing of eyeblink conditioned responses. Behav Neurosci 113, 10–18.

Meneses, A. (1999). 5-HT system and cognition. Neurosci Biobehav Rev 23, 1111–1125.

Miyazaki, K., Miyazaki, K.W., and Doya, K. (2011a). Activation of dorsal raphe serotonin neurons underlies waiting for delayed rewards. J Neurosci 31, 469–479.

Miyazaki, K., Miyazaki, K.W., and Doya, K. (2012a). The role of serotonin in the regulation of patience and impulsivity. Mol Neurobiol 45, 213–224.

Miyazaki, K., Miyazaki, K.W., Yamanaka, A., Tokuda, T., Tanaka, K.F., and Doya, K. (2018). Reward probability and timing uncertainty alter the effect of dorsal raphe serotonin neurons on patience. Nat Commun 9, 2048.

Miyazaki, K.W., Miyazaki, K., and Doya, K. (2011b). Activation of the central serotonergic system in response to delayed but not omitted rewards. Eur J Neurosci 33, 153–160.

Miyazaki, K.W., Miyazaki, K., and Doya, K. (2012b). Activation of dorsal raphe serotonin neurons is necessary for waiting for delayed rewards. J Neurosci 32, 10451–10457.

Miyazaki, K.W., Miyazaki, K., Tanaka, K.F., Yamanaka, A., Takahashi, A., Tabuchi, S., and Doya, K. (2014). Optogenetic activation of dorsal raphe serotonin neurons enhances patience for future rewards. Curr Biol 24, 2033–2040.

Nagai, Y., Miyakawa, N., Takuwa, H., Hori, Y., Oyama, K., Ji, B., Takahashi, M., Huang, X.P., Slocum, S.T., DiBerto, J.F., et al. (2020). Deschloroclozapine, a potent and selective chemogenetic actuator enables rapid neuronal and behavioral modulations in mice and monkeys. Nat Neurosci 23, 1157–1167.

Oristaglio, J., Hyman West, S., Ghaffari, M., Lech, M.S., Verma, B.R., Harvey, J.A., Welsh, J.P., and Malone, R.P. (2013). Children with autism spectrum disorders show abnormal conditioned response timing on delay, but not trace, eyeblink conditioning. Neuroscience 248, 708–718.

Owald, D., Felsenberg, J., Talbot, C.B., Das, G., Perisse, E., Huetteroth, W., and Waddell, S. (2015). Activity of defined mushroom body output neurons underlies learned olfactory behavior in Drosophila. Neuron 86, 417–427.

Papadopoulou, M., Cassenaer, S., Nowotny, T., and Laurent, G. (2011). Normalization for sparse encoding of odors by a wide-field interneuron. Science 332, 721–725.

Park, S.B., Coull, J.T., McShane, R.H., Young, A.H., Sahakian, B.J., Robbins, T.W., and Cowen, P.J. (1994). Tryptophan depletion in normal volunteers produces selective impairments in learning and memory. Neuropharmacology 33, 575–588.

Pavlov, I.P., and Anrep, G.V. (1927). Conditioned reflexes; an investigation of the physiological activity of the cerebral cortex (London: Oxford University Press: Humphrey Milford).

Pawlak, V., Wickens, J.R., Kirkwood, A., and Kerr, J.N. (2010). Timing is not Everything: Neuromodulation Opens the STDP Gate. Front Synaptic Neurosci 2, 146.

Perisse, E., Owald, D., Barnstedt, O., Talbot, C.B., Huetteroth, W., and Waddell, S. (2016). Aversive Learning and Appetitive Motivation Toggle Feed-Forward Inhibition in the Drosophila Mushroom Body. Neuron 90, 1086–1099.

Perrett, S.P., Ruiz, B.P., and Mauk, M.D. (1993). Cerebellar cortex lesions disrupt learning- dependent timing of conditioned eyelid responses. J Neurosci 13, 1708–1718.

Placais, P.Y., Trannoy, S., Friedrich, A.B., Tanimoto, H., and Preat, T. (2013). Two pairs of mushroom body efferent neurons are required for appetitive long-term memory retrieval in Drosophila. Cell Rep 5, 769–780.

Qian, Y., Cao, Y., Deng, B., Yang, G., Li, J., Xu, R., Zhang, D., Huang, J., and Rao, Y. (2017). Sleep homeostasis regulated by 5HT2b receptor in a small subset of neurons in the dorsal fan- shaped body of drosophila. Elife 6.

Qin, H., Cressy, M., Li, W., Coravos, J.S., Izzi, S.A., and Dubnau, J. (2012). Gamma neurons mediate dopaminergic input during aversive olfactory memory formation in Drosophila. Curr Biol 22, 608–614.

Ren, J., Friedmann, D., Xiong, J., Liu, C.D., Ferguson, B.R., Weerakkody, T., DeLoach, K.E., Ran, C., Pun, A., Sun, Y., et al. (2018). Anatomically Defined and Functionally Distinct Dorsal Raphe Serotonin Sub-systems. Cell 175, 472–487 e420.

Ries, A.S., Hermanns, T., Poeck, B., and Strauss, R. (2017). Serotonin modulates a depression- like state in Drosophila responsive to lithium treatment. Nat Commun 8, 15738.

Roth, B.L. (2016). DREADDs for Neuroscientists. Neuron 89, 683–694.

Sabandal, J.M., Berry, J.A., and Davis, R.L. (2021). Dopamine-based mechanism for transient forgetting. Nature.

Saudou, F., Boschert, U., Amlaiky, N., Plassat, J.L., and Hen, R. (1992). A family of Drosophila serotonin receptors with distinct intracellular signalling properties and expression patterns. EMBO J 11, 7–17.

Scheffer, L.K., Xu, C.S., Januszewski, M., Lu, Z., Takemura, S.Y., Hayworth, K.J., Huang, G.B., Shinomiya, K., Maitlin-Shepard, J., Berg, S., et al. (2020). A connectome and analysis of the adult Drosophila central brain. Elife 9.

Scheunemann, L., Placais, P.Y., Dromard, Y., Schwarzel, M., and Preat, T. (2018). Dunce Phosphodiesterase Acts as a Checkpoint for Drosophila Long-Term Memory in a Pair of Serotonergic Neurons. Neuron 98, 350–365 e355.

Schroll, C., Riemensperger, T., Bucher, D., Ehmer, J., Voller, T., Erbguth, K., Gerber, B., Hendel, T., Nagel, G., Buchner, E., et al. (2006). Light-induced activation of distinct modulatory neurons triggers appetitive or aversive learning in Drosophila larvae. Curr Biol 16, 1741–1747.

Schwaerzel, M., Monastirioti, M., Scholz, H., Friggi-Grelin, F., Birman, S., and Heisenberg, M. (2003). Dopamine and octopamine differentiate between aversive and appetitive olfactory memories in Drosophila. J Neurosci 23, 10495–10502.

Sejourne, J., Placais, P.Y., Aso, Y., Siwanowicz, I., Trannoy, S., Thoma, V., Tedjakumala, S.R., Rubin, G.M., Tchenio, P., Ito, K., et al. (2011). Mushroom body efferent neurons responsible for aversive olfactory memory retrieval in Drosophila. Nat Neurosci 14, 903–910.

Shuai, Y., Hu, Y., Qin, H., Campbell, R.A., and Zhong, Y. (2011). Distinct molecular underpinnings of Drosophila olfactory trace conditioning. Proc Natl Acad Sci U S A 108, 20201–20206.

Sitaraman, D., Zars, M., Laferriere, H., Chen, Y.C., Sable-Smith, A., Kitamoto, T., Rottinghaus, G.E., and Zars, T. (2008). Serotonin is necessary for place memory in Drosophila. Proc Natl Acad Sci U S A 105, 5579–5584.

Skinner, B.F. (1938). The behavior of organisms (New York,: Appleton-Century-Crofts).

Stahl, A., Noyes, N.C., Boto, T., Jing, M., Zeng, J., King, L.B., Li, Y., Davis, R.L., and Tomchik, S.M. (2021). Associative learning drives longitudinally-graded presynaptic plasticity of neurotransmitter release along axonal compartments. bioRxiv.

Suzuki, Y., Schenk, J.E., Tan, H., and Gaudry, Q. (2020). A Population of Interneurons Signals Changes in the Basal Concentration of Serotonin and Mediates Gain Control in the Drosophila Antennal Lobe. Curr Biol 30, 1110–1118 e1114.

Takemura, S.Y., Aso, Y., Hige, T., Wong, A., Lu, Z., Xu, C.S., Rivlin, P.K., Hess, H., Zhao, T., Parag, T., et al. (2017). A connectome of a learning and memory center in the adult Drosophila brain. Elife 6.

Tanaka, N.K., Tanimoto, H., and Ito, K. (2008). Neuronal assemblies of the Drosophila mushroom body. J Comp Neurol 508, 711–755.

Tanimoto, H., Heisenberg, M., and Gerber, B. (2004). Experimental psychology: event timing turns punishment to reward. Nature 430, 983.

Thornquist, S.C., Langer, K., Zhang, S.X., Rogulja, D., and Crickmore, M.A. (2020). CaMKII Measures the Passage of Time to Coordinate Behavior and Motivational State. Neuron 105, 334–345 e339.

Thornquist, S.C., Pitsch, M.J., Auth, C.S., and Crickmore, M.A. (2021). Biochemical evidence accumulates across neurons to drive a network-level eruption. Mol Cell 81, 675–690 e678.

Tomchik, S.M., and Davis, R.L. (2009). Dynamics of learning-related cAMP signaling and stimulus integration in the Drosophila olfactory pathway. Neuron 64, 510–521.

Tully, T., and Quinn, W.G. (1985). Classical conditioning and retention in normal and mutant Drosophila melanogaster. J Comp Physiol A 157, 263–277.

Turner, G.C., Bazhenov, M., and Laurent, G. (2008). Olfactory representations by Drosophila mushroom body neurons. J Neurophysiol 99, 734–746.

Waddell, S. (2016). Neural Plasticity: Dopamine Tunes the Mushroom Body Output Network. Curr Biol 26, R109–112.

Waddell, S., Armstrong, J.D., Kitamoto, T., Kaiser, K., and Quinn, W.G. (2000). The amnesiac gene product is expressed in two neurons in the Drosophila brain that are critical for memory. Cell 103, 805–813.

Wan, J., Peng, W., Li, X., Qian, T., Song, K., Zeng, J., Deng, F., Hao, S., Feng, J., Zhang, P., et al. (2021). A genetically encoded sensor for measuring serotonin dynamics. Nat Neurosci.

Wang, Y., Mamiya, A., Chiang, A.S., and Zhong, Y. (2008). Imaging of an early memory trace in the Drosophila mushroom body. J Neurosci 28, 4368–4376.

Wittmann, M., Carter, O., Hasler, F., Cahn, B.R., Grimberg, U., Spring, P., Hell, D., Flohr, H., and Vollenweider, F.X. (2007). Effects of psilocybin on time perception and temporal control of behaviour in humans. J Psychopharmacol 21, 50–64.

Woodruff-Pak, D.S., and Papka, M. (1996). Huntington’s disease and eyeblink classical conditioning: normal learning but abnormal timing. J Int Neuropsychol Soc 2, 323–334.

Wu, C.L., Shih, M.F., Lai, J.S., Yang, H.T., Turner, G.C., Chen, L., and Chiang, A.S. (2011). Heterotypic gap junctions between two neurons in the drosophila brain are critical for memory. Curr Biol 21, 848–854.

Wu, Y., Ren, Q., Li, H., and Guo, A. (2012). The GABAergic anterior paired lateral neurons facilitate olfactory reversal learning in Drosophila. Learn Mem 19, 478–486.

Yu, D., Akalal, D.B., and Davis, R.L. (2006). Drosophila alpha/beta mushroom body neurons form a branch-specific, long-term cellular memory trace after spaced olfactory conditioning. Neuron 52, 845–855.

Yu, D., Keene, A.C., Srivatsan, A., Waddell, S., and Davis, R.L. (2005). Drosophila DPM neurons form a delayed and branch-specific memory trace after olfactory classical conditioning. Cell 123, 945–957.

Yuan, Q., Lin, F., Zheng, X., and Sehgal, A. (2005). Serotonin modulates circadian entrainment in Drosophila. Neuron 47, 115–127.

Zhang, S., and Roman, G. (2013). Presynaptic inhibition of gamma lobe neurons is required for olfactory learning in Drosophila. Curr Biol 23, 2519–2527.

Zhang, X., Coates, K., Dacks, A., Gunay, C., Lauritzen, J.S., Li, F., Calle-Schuler, S.A., Bock, D., and Gaudry, Q. (2019a). Local synaptic inputs support opposing, network-specific odor representations in a widely projecting modulatory neuron. Elife 8.

Zhang, X., Noyes, N.C., Zeng, J., Li, Y., and Davis, R.L. (2019b). Aversive Training Induces Both Presynaptic and Postsynaptic Suppression in Drosophila. J Neurosci 39, 9164–9172.

Zhang, Y., Lu, H., and Bargmann, C.I. (2005). Pathogenic bacteria induce aversive olfactory learning in Caenorhabditis elegans. Nature 438, 179–184.

Zhou, M., Chen, N., Tian, J., Zeng, J., Zhang, Y., Zhang, X., Guo, J., Sun, J., Li, Y., Guo, A., et al. (2019). Suppression of GABAergic neurons through D2-like receptor secures efficient conditioning in Drosophila aversive olfactory learning. Proc Natl Acad Sci U S A 116, 5118–5125.

